# FXR-mediated recruitment of PPP1CB suppresses SMAD2/3 phosphorylation to mitigate pulmonary fibrosis

**DOI:** 10.64898/2026.07.24.740478

**Authors:** Wenqi Li, Qiong Wu, Yongzhi Lu, Xiaoqin Li, Miao Lei, Yi Xia, Xianjie Qiu, Miru Tang, Zheng Li, Yang Peng, Wenxiang Hu, Wei K. Zhang, Jie Zheng, Cong Ma, Jinsai Shang

## Abstract

Idiopathic pulmonary fibrosis (IPF) is a progressive and fatal interstitial lung disease with limited treatment options and poorly understood molecular underpinnings. Dysregulated TGF-β/SMAD signaling is a key driver of fibrotic remodeling, promoting persistent myofibroblast activation and excessive extracellular matrix deposition. Here we identify the Farnesoid X receptor (FXR), a bile acid activated nuclear receptor, as a previously unrecognized suppressor of pulmonary fibrosis. FXR expression is significantly reduced in lung tissues from patients with PF and in myofibroblasts derived from BLM-induced mouse models, correlating inversely with fibrosis severity. Genetic ablation of FXR exacerbates BLM-induced pulmonary fibrosis by promoting fibroblast hyperactivation and dysregulation of the TGF-β/SMAD signaling pathway. Mechanistically, we identify PPP1CB as a previously unrecognized FXR-interacting protein in primary myofibroblasts derived from IPF patients. We further show that FXR both increases chromatin accessibility at the PPP1CB locus and assembles a functional complex with PPP1CB, which in turn promotes SMAD2/3 dephosphorylation and suppresses their nuclear translocation. Notably, the clinical-stage FXR agonist TERN101 exhibits potent therapeutic efficacy in a BLM-induced mouse pulmonary fibrosis model. These findings establish FXR as a critical antifibrotic regulator in lung tissue and suggest that pharmacological activation of FXR may offer a promising therapeutic strategy for IPF.

## Introduction

Idiopathic pulmonary fibrosis (IPF) is a chronic, progressive and fatal interstitial lung disease with unclear etiology, marked by recurrent alveolar epithelial injury, aberrant fibroblast proliferation and differentiation into contractile myofibroblast, as well as overwhelming extracellular matrix (ECM) accumulation in the lung interstitium^1–3^. Activated fibroblasts are the key effector cells in the progression of IPF. They exhibit strong contractile ability and produce large amounts of collagen, leading to tissue destruction and aberrant lung repair^4–6^. IPF is associated with a poor prognosis, with a median survival of only 3-5 years following diagnosis^4^. Although the advent of antifibrotic therapies such as pirfenidone and nintedanib has improved symptom management and slowed disease progression, the overall incidence and mortality rate of IPF remain largely unchanged^7–9^. Therefore, elucidating the pathogenesis of IPF and identifying novel therapeutic strategies remain urgent priorities.

The transforming growth factor-β (TGF-β) family represents the largest group of cytokines, with TGF-β acts as a potent fibrogenic signal that stimulates fibroblasts, connective tissue cells, and epithelial cells to produce and remodel the ECM^10–12^. These TGF-β family members signal through two transmembrane receptors, type I and type II TGF-β receptors, with intracellular SMAD proteins acting as key mediators of the downstream signaling responses^13,14^. Upon ligand binding, the TGF-β receptors are activated and phosphorylate regulatory Smad (R-Smad) proteins, specifically SMAD2 or SMAD3. The phosphorylated R-SMADs then form complexes with SMAD4 and translocate to the nucleus, where they regulate the transcription of diverse target genes^15,16^. Aberrant activation of this pathway leads to excessive secretion of pro-fibrotic factors, including collagen, fibronectin (FN1), and α-SMA (also known as ACTA2)^17,18^. An increasing accumulation of evidence suggests that the TGF-β/SMAD signaling pathway plays a crucial part in the pathogenesis of pulmonary fibrosis. Notably, SMAD2/3 activation exhibits a direct dependence on TGF-β stimulation, with its downstream targets significantly enriched in fibrotic core processes such as ECM deposition and myofibroblast activation^19,20^. These findings imply that selective inhibition of SMAD2/3 may offer enhanced therapeutic specificity for fibrotic disorders.

Farnesoid X receptor (FXR), encoded by the NR1H4 gene, is a bile acid-activated nuclear receptor belonging to the ligand-dependent transcription factor superfamily^21,22^. Compelling evidence establishes FXR as a master regulator of metabolic homeostasis, bile acid metabolism, and the functional integrity of multiple organs^23^. While FXR has been well-characterized as a potent anti-fibrotic target in classical organs (such as liver^24^, kidney^25^, and intestine^26^), emerging evidence highlights its critical physiological roles in non-canonical tissues, including the vascular ^27^ and pulmonary systems^28^. Notably, the FXR agonist obeticholic acid (OCA) demonstrates potent anti-inflammatory effects and ameliorates tissue remodeling in monocrotaline (MCT)-induced pulmonary arterial hypertension models^29^. Targeting FXR reduces susceptibility to SARS-CoV-2 infection in vitro, in vivo and in ex vivo–perfused human lungs and livers^30^. Although the above studies suggest a potential protective role of FXR in pulmonary pathological processes, the precise molecular mechanisms underlying its involvement in pulmonary fibrosis remain largely unclear.

Our study reveals that FXR expression is significantly downregulated in lung tissues from patients with pulmonary fibrosis, with expression levels inversely correlating with disease severity. Furthermore, genetic ablation of FXR exacerbates BLM-induced pulmonary fibrosis by promoting fibroblast hyperactivation and dysregulation of the TGF-β/SMAD signaling pathway. Mechanistically, we identify PPP1CB as a previously unrecognized FXR-interacting protein in primary myofibroblasts derived from IPF patients. We show that FXR recruits PPP1CB to dephosphorylate SMAD2/3, promoting its nuclear export and thereby inhibiting myofibroblast activation and downstream profibrotic signaling. Here we demonstrate that TERN101, a clinical-stage nanomolar nonsteroidal FXR agonist^31^, exerts robust therapeutic effects in a mouse model of pulmonary fibrosis. Collectively, these findings identify FXR as a central regulator of pulmonary fibrosis and reveal its potential as a previously unexplored therapeutic target.

## Results

### Decreased expression of FXR is correlated with disease severity of pulmonary fibrosis

To elucidate the role of FXR in pulmonary fibrosis, we first analyzed publicly available microarray datasets, which revealed a significant downregulation of *FXR* expression in lung tissues of IPF patients compared with healthy controls (**Fig. 1a**). Subsequent correlation analysis revealed that the expression of FXR was negatively correlated with the marker genes of collagen deposition (*COL3A1*), myofibroblast activation (*ACTA2*), and extracellular matrix deposition (*MMP9*) (**Fig. 1b-d**). Clinically, FXR expression was positively correlated with pulmonary function parameters (DLCO, FEV_1_, FVC) in IPF patients (**Supplementary Fig. 1a**), but showed no such correlation in healthy individuals (**Supplementary Fig. 1b**). Our analysis of single-cell transcriptomic data from patients with IPF revealed that, compared with healthy controls, *FXR* expression was reduced in fibroblasts in IPF (**Supplementary Fig. 2a-b**). To validate the scRNA-seq findings, we further performed qPCR analysis on lung tissue samples from healthy donors and patients with IPF. The results showed that *FXR* mRNA expression was significantly decreased in IPF lungs, whereas the expression of fibrosis-related genes was markedly increased (**Fig. 1e**). Immunohistochemical (IHC) staining further demonstrated that FXR expression was substantially lower in lung tissues from IPF patients than in healthy controls (**Fig. 1f**).

**Fig. 1.**
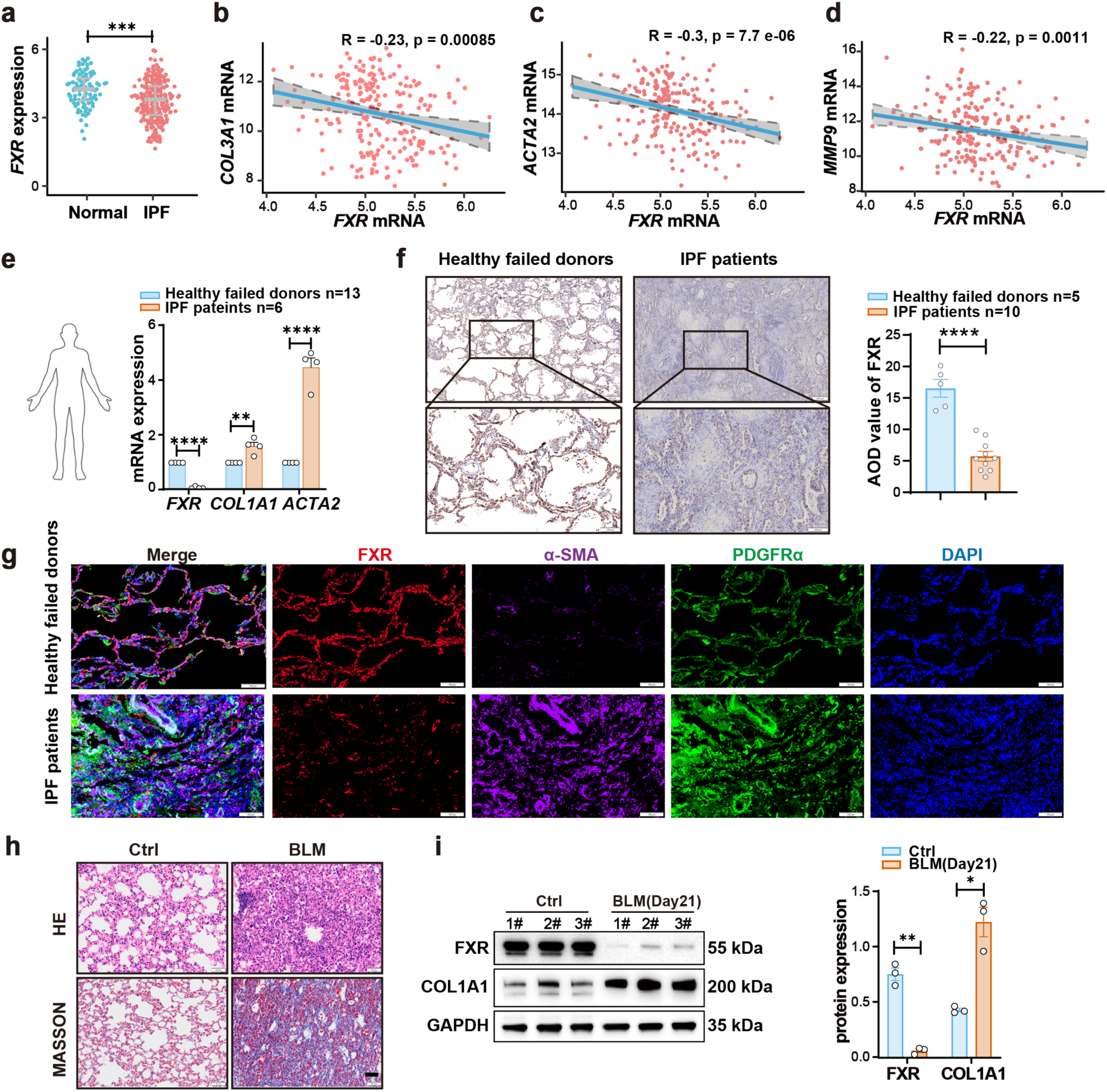
Decreased expression of FXR is correlated with disease severity of pulmonary fibrosis. (**a**) Scatterplots of FXR expression in lung tissues from public microarray datasets. (GEO: GSE47460; ILD [interstitial lung disease]). (**b-d**) Correlation analysis between the mRNA levels of lung fibrosis marker *COL3A1, α-SMA, MMP9* and *FXR* mRNA level in lung tissues from patients with IPF. (GEO: GSE47460, Spearman correlation analysis). (**e**) qPCR analysis of FXR mRNA expression in lung tissue from healthy donors (n=10) and IPF patients (n=6). (**f**) IHC staining for FXR in lung tissues from IPF patients (n=10) and healthy controls (n=5). Scale bar, 200 μm and 100 μm. (**g**) Multiplex immunofluorescence staining for FXR, α-SMA and PDGFRα in lung tissues from IPF patients and healthy controls. Scale bars, 100 μm. (**h**) HE staining and Masson staining of fibrotic lung tissues. Scale bars, 50 μm. (**i**) The protein expression of FXR and COL1A1 in mouse primary myofibroblasts. The data (e, f and i) were assessed by two-tailed Student’s t test. *\*p* < 0.05; *\*\*p* < 0.01; *\*\*\*p* < 0.001; *\*\*\*\*p* < 0.0001; ns, not significant. Data were presented as mean ± SEM. GAPDH levels were used as an internal normalization control.

To define the major cell types expressing FXR, we further analysed the IPF scRNA-seq dataset and found that *FXR* was predominantly enriched in myofibroblasts and smooth muscle cells (**Supplementary Fig. 2c**). In the scRNA-seq from the BLM-induced mouse model of pulmonary fibrosis, *Fxr* was likewise mainly expressed in mesothelial cells and smooth muscle cells (SM) (**Supplementary Fig. 2d**). To validate these localization patterns, we performed multiplex immunofluorescence staining on lung tissue sections from IPF patients and healthy controls. The results showed that α-SMA signals were markedly increased, whereas FXR signals were clearly reduced in IPF samples. Further co-localization analysis with the fibroblast marker PDGFRα demonstrated that FXR expression was detectable in the PDGFRα+ fibroblast population (**Fig. 1g, Supplementary Fig.2e**).

In the widely used BLM-induced mouse model of pulmonary fibrosis^32^, BLM-treated mice exhibited weight loss (**Supplementary Fig. 2f**) and pronounced pulmonary consolidation (**Fig. 1h**). Primary myofibroblasts isolated from fibrotic lung tissues of BLM-induced mice showed significantly elevated COL1A1 expression alongside marked FXR downregulation, recapitulating the expression profile observed in IPF patients (**Fig. 1i**). Collectively, these data demonstrate a disease-specific downregulation of FXR in myofibroblasts from both IPF patients and experimental fibrosis mice models, with expression levels inversely correlating with disease severity, implicating FXR as a potential key regulator in progression of pulmonary fibrosis.

### FXR deficiency aggravates pulmonary fibrogenesis in a BLM-challenged mice model

To determine the functional contribution of FXR in pulmonary fibrosis pathogenesis, we generated Fxr knockout (*Fxr^−/−^*) mice (**Supplementary Fig. 3a**). Histopathological analysis of major organs revealed no apparent abnormalities in *Fxr^−/−^* mice (**Supplementary Fig. 3b**), indicating that the genetic ablation of FXR does not induce baseline organ pathology. Age-matched WT and *Fxr^−/−^* mice were administered BLM (3 mg/kg) via aerosolized intratracheal delivery and sacrificed on day 21 for fibrosis evaluation (**Fig. 2a**). *Fxr^−/−^* mice exhibited significantly greater body weight loss and increased mortality rates compared to WT controls over the 21-day observation period (**Fig. 2b-c**). As shown in **Fig.2d**, micro-CT imaging revealed that *Fxr^−/−^* mice exhibited more severe structural lung destruction, lung volume reduction, and extensive fibrotic lesions compared with BLM-treated WT controls at day 21 post-injury. Pulmonary function tests revealed that *Fxr^−/−^* mice developed more severe respiratory dysfunction, characterized by increased airway resistance (RI) and reduced forced vital capacity (FVC), dynamic compliance (Cchord), and functional residual capacity (FRC) compared with WT controls (**Fig. 2e**). Consistent with these results, histopathological analysis revealed exacerbated pulmonary consolidation in *Fxr^−/−^* mice, as evidenced by H&E staining, with Masson’s trichrome staining confirming significantly increased collagen deposition compared with WT controls (**Fig. 2f-g**). Immunostaining analysis revealed elevated expression of α-SMA and Ki-67 in *Fxr^−/−^* mice compared to WT controls, indicating enhanced myofibroblast activation and cellular proliferation (**Fig. 2h**).

**Fig. 2.**
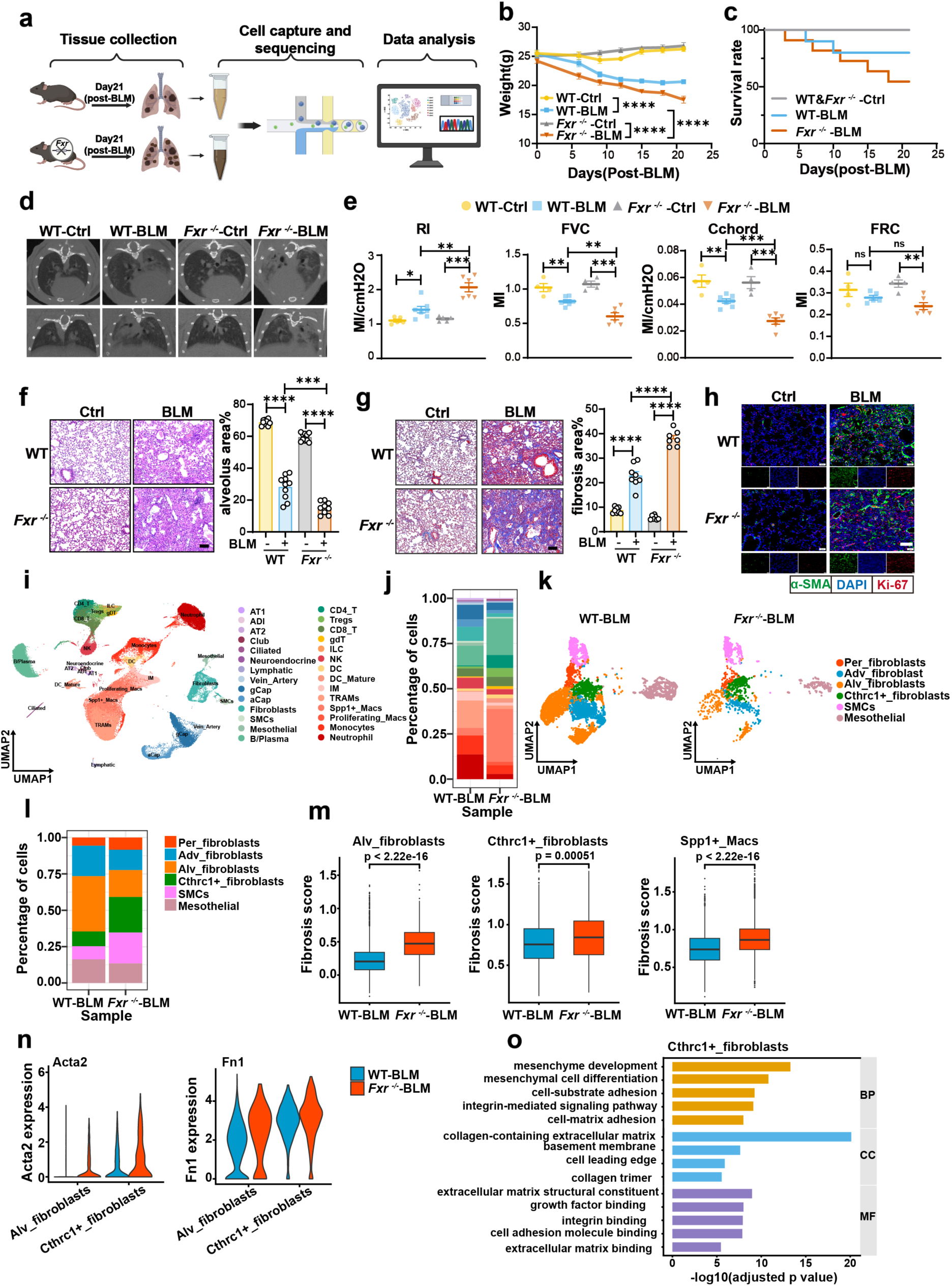
FXR deficiency accelerates the progression of pulmonary fibrosis in a BLM-challenged mouse model. (**a**) Study design for mouse lung single-cell sequencing (n=3, independent biological replicates). (**a**) created in BioRender. Wagner, D. (2025) https://BioRender.com/nmwzx11. (**b**) Body weight changes in WT and *Fxr^−/−^* mice at day 21 post-BLM instillation. (n = 5 mice per group). (**c**) Survival curves of WT and *Fxr^−/−^* mice induced at day 21 post-BLM instillation. (n = 10 mice per group). (**d**) Representative microCT images of lung tissues from WT and *Fxr^−/−^* mice with and without BLM treatment on Day 21. (**e**) Pulmonary function assessment in WT and *Fxr^−/−^* mice at day 21 post-BLM instillation. (n ≥ 5 mice per group). (**f, g**) HE staining (**f**) and Masson staining(**g**) of fibrotic areas (Scalebar:100 μm). (**h**)Immunofluorescence detection of α-SMA (green), Ki-67(red) and nuclei (DAPI, blue) in lung sections from BLM-treated mice. Scale bar: 100 μm. (**i**) UMAP of overall cell phenotypes identified from single-cell transcriptomic profiling of lungs. alveolar type (AT); Krt8+ alveolar progenitor (ADI); Venous/Artery celI (Vein_Artery); general capillary (gCAP); alveolar capillary (aCAP); Smooth Muscle Cells (SMCs); Gamma Delta T (gdT); Innate Lymphoid Cell (ILC); Natural Killer cell (NK); Dendritic Cell (DC); Interstitial macrophages (IM); Tissue-resident AM (TRAMs); Spp1+_Macs (Spp1+_Macrophage); Proliferating_Macs (Proliferating_Macrophage). (**j**) Bar plot showing the fraction of each cell type across experimental conditions. (**k**) UMAP plot of fibroblast subtypes. (**l**) Bar plot showing the fraction of fibroblasts across experimental conditions. (**m**) Bar graphs showing significant differences in fibrosis scores between the two groups. (**n**) The expression levels of Fn1 and Acta2 between the two groups. (**o**) Pathway enrichment analysis in Cthrc1+_fibroblasts. The data (b,e,f,g.m) were assessed by two-tailed Student’s t test. *\*p* < 0.05; *\*\*p* < 0.01; *\*\*\*p* < 0.001; *\*\*\*\*p* < 0.0001; ns, not significant. Data were presented as mean ± SEM.

To comprehensively characterize the alterations in cellular diversity and molecular pathways in *Fxr^−/−^* lung tissues in response to BLM, we conducted single-cell RNA sequencing (scRNA-seq) on lung tissues from BLM-treated WT and *Fxr^−/−^* mice. Following stringent quality control, normalization, and unsupervised clustering (see Methods), we identified 28 distinct cell populations (**Fig. 2i, Supplementary Fig. 3c**). Notably, the proportions of Spp1+_macrophages and proliferating_macrophages were markedly increased in *Fxr^−/−^*mice, whereas the overall fibroblast population was substantially reduced, suggesting that *Fxr^−/−^* may further exacerbate immune dysregulation and promote structural damage in lung tissue (**Fig. 2j, Supplementary Fig. 3d**). Given the central role of fibroblasts in the initiation and progression of pulmonary fibrosis^33^, we next performed a refined subcluster analysis of fibroblasts, classifying them into Peribronchial fibroblasts (Per_fibroblasts), Adventitial fibroblasts (Adv_fibroblasts), Alveolar fibroblasts (Alv_fibroblasts), and Cthrc1+ fibroblasts (Cthrc1+_fibroblasts) (**Fig. 2k, Supplementary Fig. 3e**). Cellular composition analysis revealed that, compared with WT mice, *Fxr^−/−^* mice exhibited a significant reduction in the proportion of Alv_fibroblasts, accompanied by a marked expansion of Cthrc1+_fibroblasts (**Fig. 2l, Supplementary Fig. 3f**). As Cthrc1+_fibroblasts are recognized as a highly activated pro-fibrotic fibroblast subpopulation closely associated with excessive extracellular matrix deposition and fibrosis progression^33,34^, these findings suggest that FXR deficiency promotes the transition of fibroblasts toward a more pro-fibrotic activated state. Consistently, *Fxr^−/−^* mice displayed significantly higher fibrosis scores than WT mice (**Fig. 2m**). Further gene expression analysis demonstrated that the expression levels of fibrosis-associated genes, including Acta2, Fn1, and Ltbp2, were significantly elevated in both Alv_fibroblasts and Cthrc1+_fibroblasts from *Fxr^−/−^* mice compared with WT controls (**Fig. 2n, Supplementary Fig. 3g**). In addition, pathway enrichment analysis revealed that FXR deficiency significantly upregulated fibrosis-related pathways associated with extracellular matrix remodeling and collagen deposition (**Fig. 2o, Supplementary Fig. 3h**). The collective evidences demonstrate that FXR deficiency accelerates fibrotic progression and underscore its essential role in restraining pulmonary fibrosis development.

### FXR suppresses myofibroblast activation through inhibition of SMAD2/3 phosphorylation and nuclear translocation

To systematically investigate the role of FXR in pulmonary fibrosis, we isolated and cultured primary mouse lung fibroblasts (MLFs) from both WT and *Fxr^−/−^* mice. Quantitative analysis revealed that FXR deficiency markedly potentiated TGF-β mediated expression of fibrotic genes (**Fig. 3a, Supplementary Fig. 4a**) and significantly promoted MLFs migration (**Fig. 3b, Supplementary Fig. 4b**) compared to WT controls. Conversely, FXR overexpression robustly attenuated TGF-β-induced myofibroblast activation, as evidenced by reduced expression of fibrotic genes (**Supplementary Fig. 4c**). To establish clinical relevance, we successfully isolated primary myofibroblasts from lung tissues of IPF patients, which exhibited significantly elevated expression of fibrotic markers compared to the human lung fibroblast cell line (HLF1) (**Supplementary Fig. 4d**). Notably, FXR overexpression in human lung myofibroblasts consistently recapitulated our findings, showing significant suppression of fibrotic genes expression (**Fig. 3c**). Furthermore, we employed TERN101, a clinical-stage FXR agonist with enhanced selectivity and potency, to pharmacologically activate FXR^31,35^. TERN101 treatment demonstrated a dose-dependent repression of fibrotic genes expression in both human myofibroblast (**Fig. 3d, Supplementary Fig. 4e**) and mouse fibroblast (**Fig. 3f, Supplementary Fig. 4f-g**). Under profibrotic stimulation, quiescent fibroblasts undergo phenotypic transition into activated myofibroblasts, acquiring enhanced proliferative, migratory^36^. Notably, TERN101 treatment attenuated TGF-β-induced MLFs migration (**Fig. 3e, Supplementary Fig. 4h**). In contrast, FXR antagonist DY268 and ZGG showed no significant inhibitory effects (**Supplementary Fig. 4i**). These results indicate FXR activation mediates anti-fibrotic effects by suppressing myofibroblast activation pathways.

**Fig. 3.**
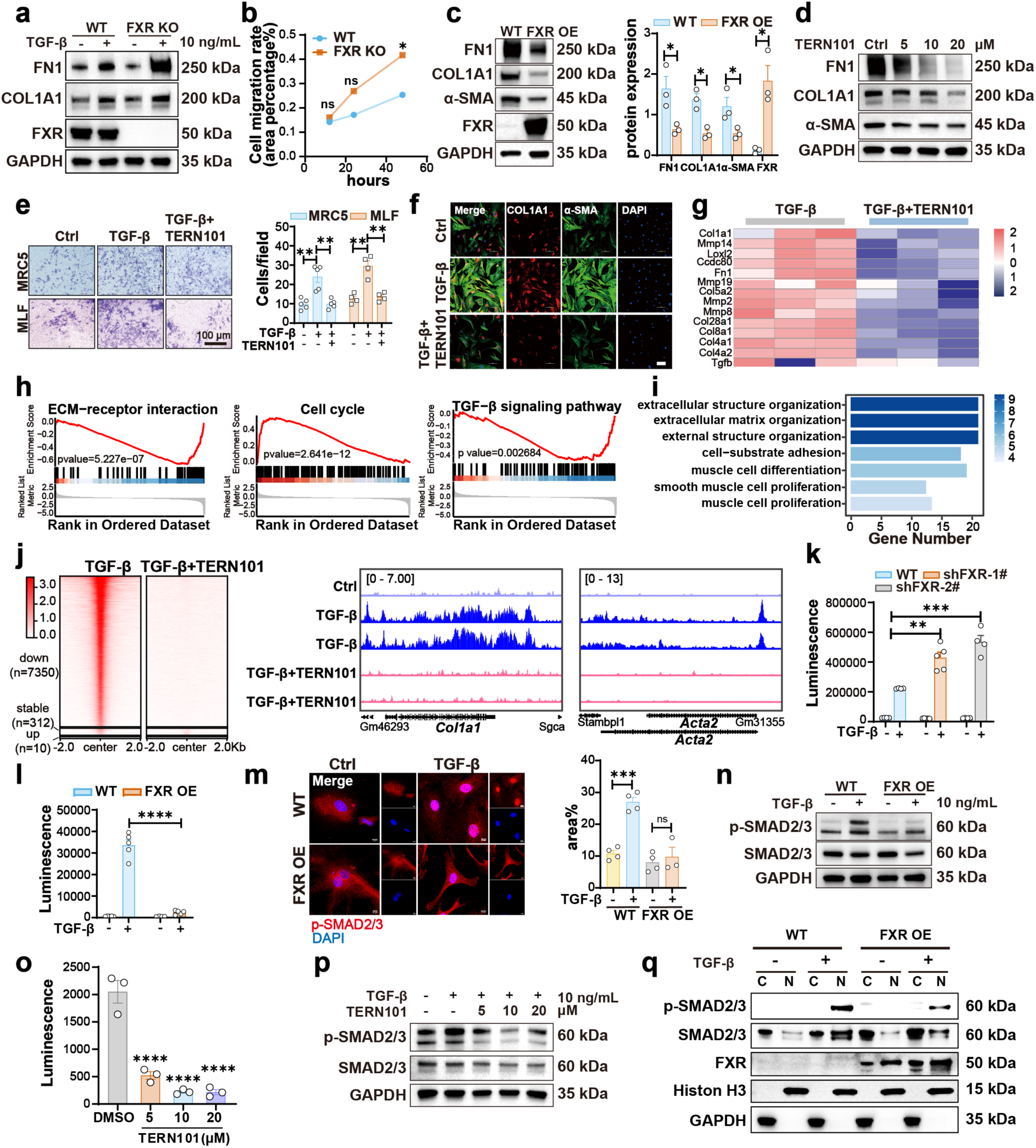
FXR suppresses myofibroblast activation through inhibition of SMAD2/3 phosphorylation and nuclear translocation. (**a**) Expression level of FN1, COL1A1 and FXR in WT and FXR KO MLFs following 48-hour TGF-β stimulation (10 ng/mL) (n=3). (**b**) Wound healing assay showing the migration of WT and FXR KO MLFs. (**c**) Western blot analysis of fibrotic genes protein levels in IPF Myofb overexpressing FXR (n=3). (**d**) Western blot analysis of FN1, COL1A1 and α-SMA expression in IPF myofb treated with TERN101 (n=3). (**e**) Transwell migration assay of MRC5 and MLFs toward 10% FBS, treated with TGF-β (10 ng/ml) and TERN101 (10 μM) for 48 h. Migrated cells were fixed, stained with crystal violet, and quantified (n ≥ 4). Scale bar: 100 μm. (**f**) Representative immunofluorescence images of MLFs treated with TGF-β (10 ng/ml) and TERN101 (10 μM) for 48 h. Scale bar: 100 μm. (**g**) A heatmap showing representative TGF-β-upregulated genes inhibited by TERN101 (n = 3 biological replicates per group). (**h**) GSEA plots showing TEN101-reversed pathways upon TGF-β stimulation. (**i**) Pathway enrichment analysis showing TGF-β-upregulated pathways inhibited by TERN101. (**j**) Heatmap showing the genomic occupancy of p-SMAD2/3 from −2 kb flanking TSS to + 2 kb flanking TES in MLFs. IGV snapshot showing the p-SMAD2/3 CUT&Tag signals at the fibrotic genes (*Col1a1*, *Acta2*) genomic loci. (**k**) Luciferase activity was detected in FXR-knockdown HEK293T-CAGA12 stable cells following stimulation with TGF-β (5 ng/mL) for 24 hours. (**l**) Luciferase activity was detected in FXR-overexpressing HEK293T-CAGA12 stable cells following stimulation with TGF-β (2 ng/mL) for 24 hours. (**m**) Representative immunofluorescence images of p-SMAD2/3 nuclear translocation in WT and FXR OE MLFs treated with or without TGF-β (10 ng/mL, 1 h). Scale bar: 10 μm. (**n**) Western blot analysis of p-SMAD2/3 and SMAD2/3 protein levels in WT and FXR OE MLFs with or without TGF-β (10 ng/mL, 1 h) stimulation (n=3). (**o**) Effects of TERN101 on SMAD-responsive CAGA12 luciferase reporter activity in HEK293T cells.(**p**) Western blot analysis of p-SMAD2/3 and SMAD2/3 protein expression in MLFs with or without TERN101(10 μM, 24h) and TGF-β (10 ng/mL, 1 h) treatment (n=4). (**q**) MRC5-FXROE or MRC5-WT cells were treated with TGFβ (10 ng/ml, 1 h). Nuclear and cytoplasmic fractions were analyzed by western blotting. The data were assessed by two-tailed Student’s t test and one-way ANOVA. *\*p* < 0.05; *\*\*p* < 0.01; *\*\*\*p* < 0.001; *\*\*\*\*p* < 0.0001; ns, not significant. Data were presented as mean ± SEM.

To comprehensively characterize transcriptomic alterations, we performed RNA sequencing on TGF-β-stimulated MLFs treated with or without TERN101. Transcriptomic analysis identified 201 upregulated and 296 downregulated genes (**Supplementary Fig. 4k**), with TERN101 treatment largely reversing TGF-β-induced gene expression alterations (**Supplementary Fig. 4l**). Notably, among all TGF-β-responsive genes, 14 fibrosis-associated genes were significantly reversed by TERN101 treatment (**Fig. 3g**). Gene set enrichment analysis (GSEA) further demonstrated that FXR activation was significantly associated with suppression of the ECM-receptor interaction, cell cycle and TGF-β signaling pathway (**Fig. 3h**). KEGG pathway enrichment analysis revealed that TERN101 downregulated signaling cascades associate with smooth muscle contraction and ECM organization (**Fig. 3i**). Strikingly, the antifibrotic effect of TERN101 was substantially attenuated in FXR-deficient cells (**Supplementary Fig. 4j**), highlighting the essential role of FXR in mediating this response. Together, these findings demonstrate that both pharmacological activation and overexpression of FXR robustly suppress myofibroblasts activation, establishing FXR as a key antifibrotic regulator in IPF.

GSEA revealed that treatment with the FXR-specific agonist TERN101 significantly suppressed the TGF-β signaling pathway (**Fig. 3h**). In the TGF-β/SMAD pathway, the receptor-regulated SMADs (R-SMADs), particularly SMAD2/3, act as central regulators of fibrotic responses by directly regulating the transcription of ECM-related genes^19,37,38^. To investigate how FXR modulates TGF-β/SMAD signaling at the chromatin level, we performed CUT&Tag profiling in MLFs using a anti-SMAD2/3-specific antibody to map genome-wide p-SMAD2/3 occupancy. TERN101 markedly reduced p-SMAD2/3 binding to the promoters of canonical fibrotic targets (*Col1a1*, *Acta2*) as revealed by CUT&Tag profiling (**Fig. 3j, Supplementary Fig. 5a**). These findings demonstrate that FXR activation attenuates fibrotic genes expression by suppressing SMAD-dependent TGF-β signal transduction. To further investigate FXR-mediated transcriptional regulation, we generated a stable HEK293T cell line expressing the SMAD2/3-responsive CAGA12-luciferase reporter construct, which exhibited robust and reproducible luminescence to TGF-β stimulation (**Supplementary Fig. 5b**). Strikingly, functional studies in HEK293T-CAGA12 reporter cells demonstrated that FXR overexpression markedly suppressed SMAD-driven transcriptional activity, whereas FXR knockdown significantly enhanced this activity. (**Fig. 3k-l, Supplementary Fig. 5c-d**). TGF-β stimulation is known to induce SMAD2/3 phosphorylation and facilitate its nuclear translocation, thereby initiating transcription of downstream target genes^39^. Immunoblotting and immunofluorescence analyses revealed that FXR overexpression reduced SMAD2/3 phosphorylation (**Fig. 3n, Supplementary Fig. 5e**) and impeded its nuclear translocation (**Fig. 3m**), whereas FXR knockout exerted the opposite effects (**Supplementary Fig. 5f-g**). To further elucidate how FXR regulates the intracellular distribution of total SMAD2/3, We performed nuclear–cytoplasmic fractionation in WT and stable FXR-overexpressing MRC5 cells. In WT cells, TGF-β stimulation robustly induced SMAD2/3 phosphorylation and promoted its nuclear accumulation. By contrast, in FXR-overexpressing cells, although TGF-β still induced SMAD2/3 phosphorylation, nuclear p-SMAD2/3 levels were markedly reduced, with a concomitant increase in their relative abundance in the cytoplasmic fraction. These findings suggest that FXR promotes dephosphorylation of nuclear p-SMAD2/3, thereby accelerating SMAD2/3 nuclear export (**Fig. 3q**). Pharmacological activation of FXR with TERN101 attenuates SMAD transcriptional activity (**Fig. 3o**), accompanied by significant reductions in SMAD2/3 phosphorylation (**Fig. 3p, Supplementary Fig. 5h)** and nuclear translocation (**Supplementary Fig. 5i**). These results demonstrate that FXR attenuates myofibroblast activation by suppressing SMAD2/3 phosphorylation and nuclear translocation, underscoring its role as a key regulator of the TGF-β/SMAD signaling axis.

### FXR-mediated recruitment of PPP1CB inhibits SMAD2/3 phosphorylation

We investigated the molecular mechanism by which FXR regulates SMAD2/3 phosphorylation by performing immunoprecipitation-mass spectrometry (IP-MS) in primary myofibroblasts derived from IPF patients (**Fig. 4a**). We identified PPP1CB as a previously unrecognized FXR-interacting protein. PPP1CB is a serine/threonine protein phosphatase that regulates diverse cellular processes through substrate-specific dephosphorylation^40^ (**Supplementary Fig. 6a**). Co-immunoprecipitation (Co-IP) assays in HEK293T cells co-transfected with FXR and PPP1CB expression plasmids confirmed the interaction between FXR and PPP1CB (**Fig. 4b**), and immunofluorescence analysis revealed their nuclear co-localization in both MLFs and A549 cells (**Fig. 4c, Supplementary Fig. 6c**). To define the FXR structural domain responsible for PPP1CB binding, we generated a series of FXR truncation mutants lacking the AF1, DBD, Hinge or LBD regions (**Fig. 4d**). Co-IP assays demonstrated that deletion of the DBD abolished the interaction with PPP1CB, whereas the other mutants retained this interaction (**Fig. 4e**). Moreover, deletion of the DBD attenuated the inhibitory effect of FXR on SMAD-mediated transcriptional activity, indicating that the DBD is required for FXR-mediated negative regulation of SMAD transcriptional output (**Supplementary Fig. 6d**). We next investigated the mechanisms by which FXR regulates downstream genes using an assay for transposase-accessible chromatin with high-throughput sequencing (ATAC-seq). ATAC-seq analysis revealed that FXR depletion reduced chromatin accessibility in MLFs, particularly at promoter regions enriched for FXR-binding motifs (**Fig. 4f**). Notably, the PPP1CB promoter exhibited diminished accessibility upon FXR knockout, indicating that FXR deficiency compromises chromatin openness at this locus (**Fig. 4j**). Gene expression analysis further demonstrated a positive correlation between FXR and PPP1CB expression levels (**Supplementary Fig. 6e-f**), and FXR overexpression enhanced their interaction (**Fig. 4h-i**). To determine whether FXR binds to the promoter of PPP1CB and enhances its transcription, we performed CUT&Tag profiling in MLFs using an FXR-specific antibody to map genome-wide FXR occupancy. The results demonstrate that FXR is enriched at the promoter region of PPP1CB in MLFs (**Supplementary Fig. 6g**). To determine whether PPP1CB affects SMAD-mediated transcription, we performed functional studies in HEK293T-CAGA12 reporter cells. PPP1CB overexpression significantly attenuated SMAD-driven transcriptional activity, whereas PPP1CB knockdown enhanced it (**Fig. 4j-k, Supplementary Fig. 6h**). Moreover, graded overexpression of PPP1CB suppressed p-SMAD2/3 levels in a dose-dependent manner (**Fig. 4l**), further supporting a negative regulatory role of PPP1CB in controlling SMAD2/3 phosphorylation. Subsequently, we performed nuclear–cytoplasmic fractionation assays in WT and stable PPP1CB-knockdown MRC5 cells. The results showed that, in WT cells, TGF-β stimulation induced redistribution of SMAD2/3 from the cytoplasm to the nucleus. In contrast, PPP1CB knockdown further enhanced TGF-β–induced accumulation of SMAD2/3 in the nuclear fraction **(Fig. 4m)**. These results further support that PPP1CB promotes SMAD2/3 dephosphorylation and thereby facilitates their nuclear export. Confocal microscopy confirmed nuclear co-localization of PPP1CB and p-SMAD2/3 in MLFs **(Supplementary Fig. 6i)**. Co-IP assays revealed an interaction between p-SMAD2/3 and PPP1CB **(Fig. 4n)**. Collectively, these data suggest that PPP1CB functions as a phosphatase for SMAD2/3, binding to p-SMAD2/3 and promoting its dephosphorylation, thereby facilitating the nuclear export of SMAD2/3. To identify catalytically important residues, we selected PPP1CB mutations reported in disease (P49R, A56P, E183A, D252Y and E274K)^41^. Notably, D252Y and E274K mutations abolished phosphatase activity, resulting in increased phosphorylation of SMAD2/3 (**Fig. 4o**). To define the structural basis, we mapped D252 and E274 onto the PPP1CB structure. Both residues lie in close proximity to the metal-ion coordination site (**Supplementary Fig. 6j**).

**Fig. 4.**
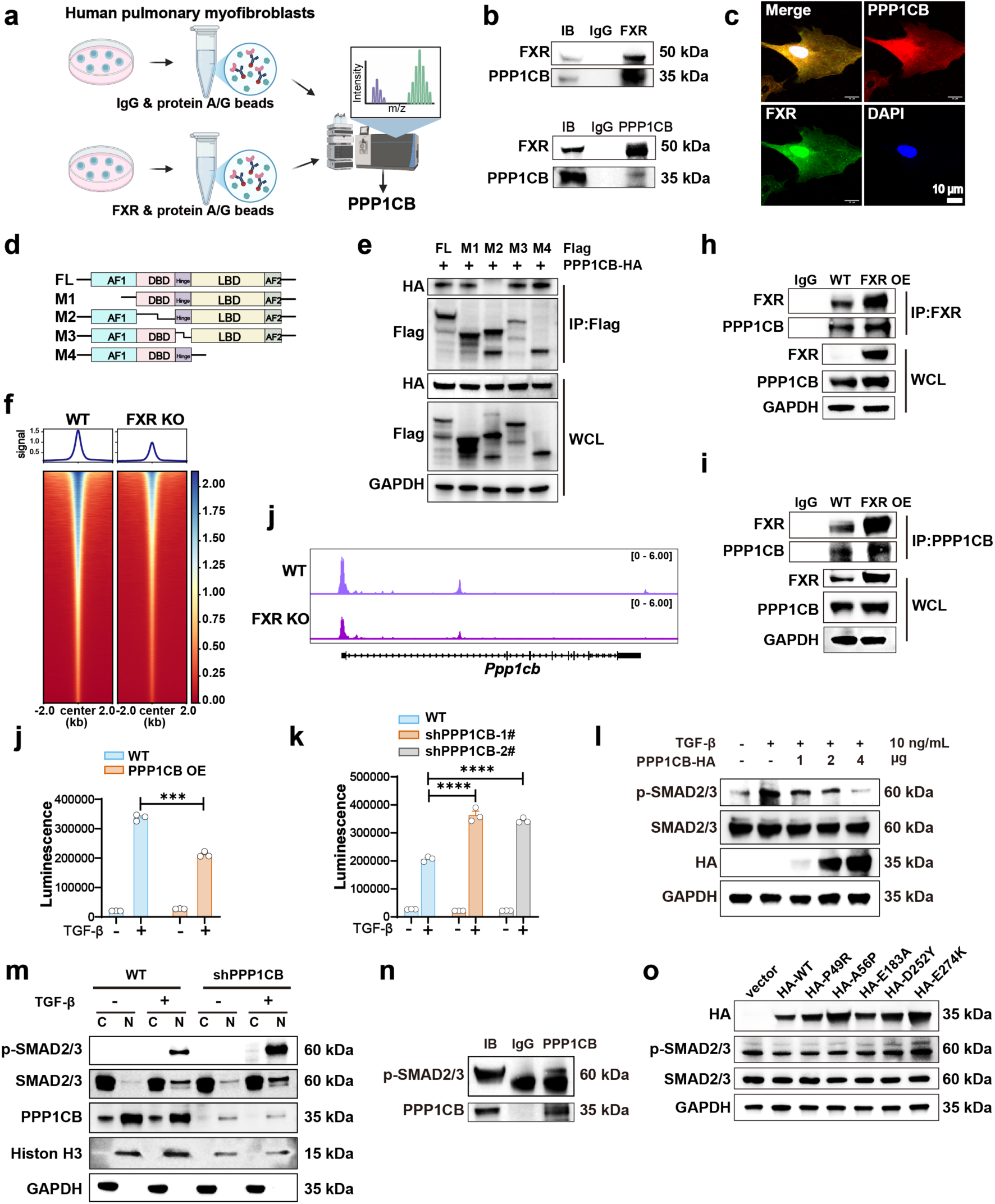
FXR-mediated recruitment of PPP1CB inhibits SMAD2/3 phosphorylation. (**a**) Immunoprecipitation (IP) of FXR from IPF myofb-FXROE cells followed by LC-MS/MS analysis to identify interacting proteins. The schematic(**a**) was created in BioRender. Wagner, D. (2025) https://BioRender.com/nmwzx11. (**b**) CO-IP analysis of the interaction between FXR and PPP1CB in HEK293T (n=3). (**c**) Immunofluorescence localization of PPP1CB and FXR in MLFs. Scale bar: 10 μm. (**d**) Schematic illustrations of FXR truncation plasmids. (**e**) Immunoblot analysis following cotransfection with PPP1CB-FL and FXR truncation plasmids. (**f**) Heatmap of chromatin accessibility analysis by ATAC-seq in WT and FXR KO MLFs. (**j**) IGV analysis of ATAC-seq signals in the gene locus of *Ppp1cb* in WT and FXR KO MLFs. (**h-i**) IP analysis of the interaction between FXR and PPP1CB in A549 cells overexpressing FXR. (**j-k**) Luciferase reporter assays in HEK293T cells stably expressing the SMAD-responsive CAGA12-luciferase with PPP1CB overexpression(**j**) or knockdown(**k**) (n=3). (**l**) HEK293T cells were transfected with increasing doses of HA-PPP1CB, stimulated with TGF-β (10 ng/mL) for 1 h, and analyzed by immunoblotting with the indicated antibodies. (**m**) MRC5-shPPP1CB or MRC5-WT cells were treated with TGFβ (10 ng/ml, 1 h). Nuclear and cytoplasmic fractions were analyzed by western blotting. (**n**) CO-IP analysis of the interaction between p-SMAD2/3 and PPP1CB in HEK293T. (**o**) Detection of SMAD2/3 phosphorylation in HEK293T cells transfected with PPP1CB-WT or PPP1CB mutation plasmids. The data were assessed by two-tailed Student’s t test. *\*p* < 0.05; *\*\*p* < 0.01; *\*\*\*p* < 0.001; *\*\*\*\*p* < 0.0001; ns, not significant. Data were presented as mean ± SEM.

Based on these observations, we hypothesized that FXR recruits PPP1CB to mediate SMAD2/3 dephosphorylation, thereby attenuating transcription of downstream fibrotic genes. Co-overexpression of FXR and PPP1CB synergistically inhibited SMAD transcriptional activity to a greater extent than alone (**Fig. 5a**). Consistently, western blot analysis showed that co-expression of FXR and PPP1CB produced a stronger reduction in SMAD2/3 phosphorylation than PPP1CB overexpression alone, indicating a cooperative role for FXR and PPP1CB in the negative regulation of SMAD signaling (**Fig. 5b**). To determine whether FXR-mediated inhibition of SMAD2/3 phosphorylation is dependent on PPP1CB, we found that the FXR agonist TERN101 attenuated SMAD2/3 phosphorylation in MLFs, an effect reversed by co-treatment with the PPP1CB inhibitor okadaic acid (**Fig. 5c**). To exclude potential off-target effects of okadaic acid on other PPP1 isoforms, we generated stable cell lines with FXR overexpression and PPP1CB knockdown, confirming that PPP1CB depletion abrogates FXR-mediated SMAD2/3 dephosphorylation (**Fig. 5d**). Finally, to assess whether SMAD2/3 reciprocally influences the formation of the FXR–PPP1CB complex, we established stable SMAD2/3 knockdown cell lines and observed that depletion of SMAD2/3 did not affect the interaction between FXR and PPP1CB, indicate that the interaction between FXR and PPP1CB is independent of SMAD2/3 (**Fig. 5e-f**). Collectively, our data demonstrate that FXR recruits PPP1CB to dephosphorylate SMAD2/3, thereby attenuating myofibroblast activation and exerting antifibrotic effects.

**Fig. 5.**
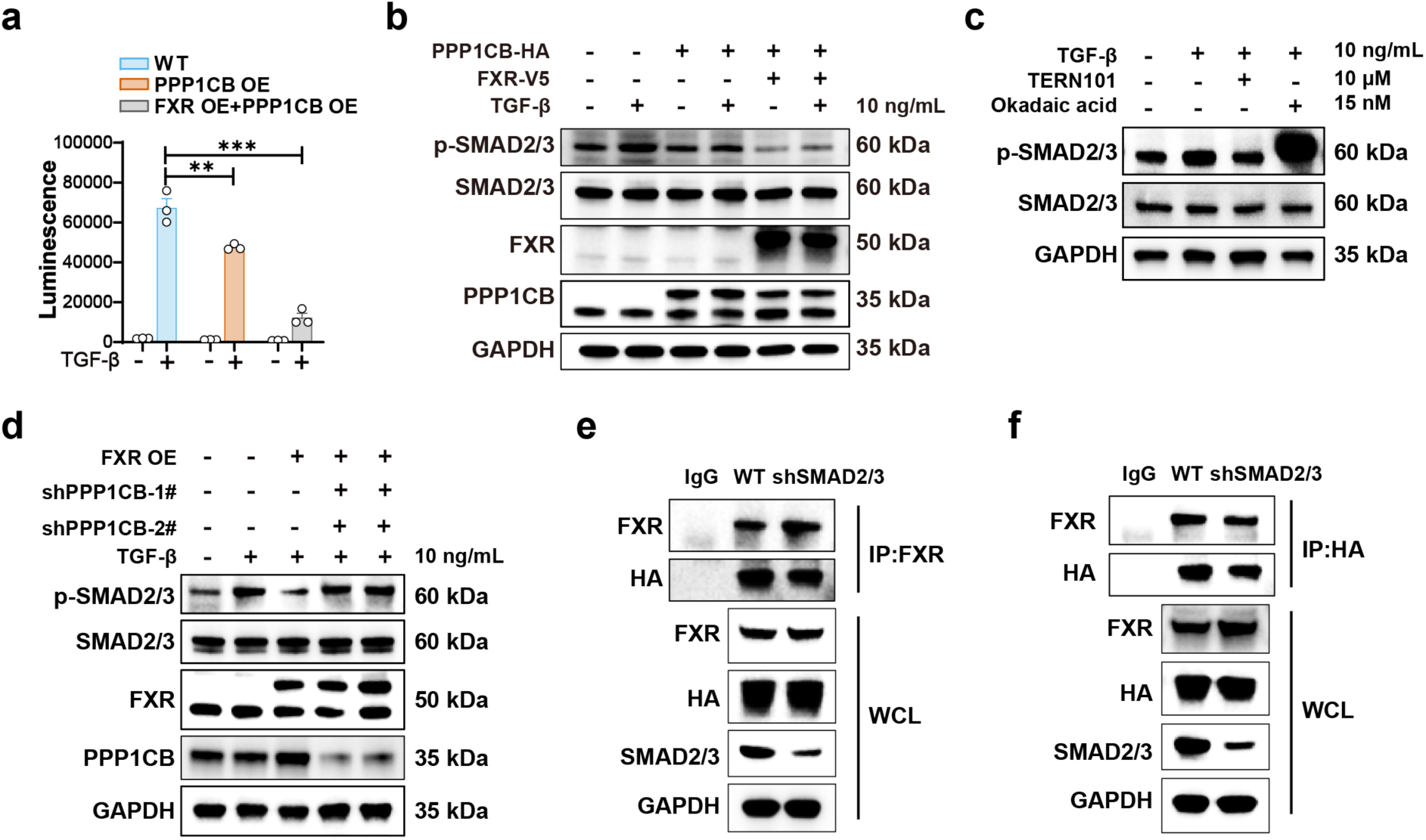
FXR-mediated recruitment of PPP1CB inhibits SMAD2/3 phosphorylation. **(a)** Luciferase reporter assays in HEK293T cells stably expressing the SMAD-responsive CAGA12-luciferase with the PPP1CB and FXR overexpression (TGF-β=5ng/mL) (n=3). **(b)** HEK293T cells transfected with PPP1CB alone or with PPP1CB and FXR were stimulated with or without TGF-β for 1 hour, following which SMAD2/3 phosphorylation levels were assessed. **(c)** In MLFs, following treatment with TERN101 for 48 hours and Okadaic acid for 24 hours, the phosphorylation levels of SMAD2/3 were detected after subsequent stimulation with TGF-β for 1 hour. **(d)** SMAD2/3 phosphorylation was detected in A549 cells with FXR overexpression and PPP1CB knockdown following stimulation with TGF-β for 1 hour. (**e-f**) CO-IP analysis of the interaction between FXR and PPP1CB in WT and SMAD2/3-knockdown HEK293T cells. The data were assessed by two-tailed Student’s t test. *\*p* < 0.05; *\*\*p* < 0.01; *\*\*\*p* < 0.001; *\*\*\*\*p* < 0.0001; ns, not significant. Data were presented as mean ± SEM.

### Pharmacological activation of FXR attenuates pulmonary fibrosis progression in mice

Building upon our mechanistic findings, we next evaluated the therapeutic potential of FXR activation in mice models of pulmonary fibrosis. FXR agonists have shown promising clinical efficacy in liver diseases including primary biliary cholangitis (PBC), primary sclerosing cholangitis (PSC), and non-alcoholic steatohepatitis (NASH)^42^. We selected three pharmacological agonists for comprehensive characterization: OCA (steroidal), GW4064 (selective non-steroidal) and TERN101(partial agonist). TR-FRET assays showed all three compounds induced dose-dependent coactivator recruitment of the SRC2-2 coactivator peptide to FXR (EC_50_: OCA, 88 nM; GW4064, 46 nM, TERN101, 57 nM) (**Supplementary Fig. 7a**). While luciferase reporter assays revealed that TERN101 elicited the more potent FXR-mediated transcriptional activation than OCA (EC_50_: OCA, 937.2 nM, GW4064, 76 nM, TERN101, 79.6 nM) (**Supplementary Fig. 7b**). ITC further confirmed the highest binding affinity of TERN101 among the three compounds (**Fig. 6q, Supplementary Fig. 7c**). To validate binding specificity, we solved the crystal structure of FXR-LBD in complex with TERN101 at 2.95 Å resolution, which revealed canonical agonist interactions (**Fig. 6r, Supplementary Fig. 10**). Moreover, HDX-MS analysis showed a greater reduction in deuterium uptake upon TERN101 binding compared to OCA and GW4064, suggesting superior conformational stabilization of FXR with in the ligand binding pocket (**Supplementary Fig. 7g, Supplementary Fig. 8**). We next assessed the anti-fibrotic efficacy of the three FXR agonists in both human and mice fibroblasts. The results revealed that all compounds significantly downregulated the mRNA and protein levels of the key fibrotic genes. Among them, TERN101 exhibited the most pronounced inhibitory effects (**Supplementary Fig. 7d-e**). Moreover, cell viability assays further demonstrated that OCA and TERN101 had significantly lower cytotoxicity compared to GW4064 (**Supplementary Fig. 7f**). OCA has been reported to have pruritus as a side effect^43^. Taken together, these findings identify TERN101 as the most suitable FXR agonist for further validation in mice models, based on its superior anti-fibrotic potency and minimal cytotoxicity.

**Fig. 6.**
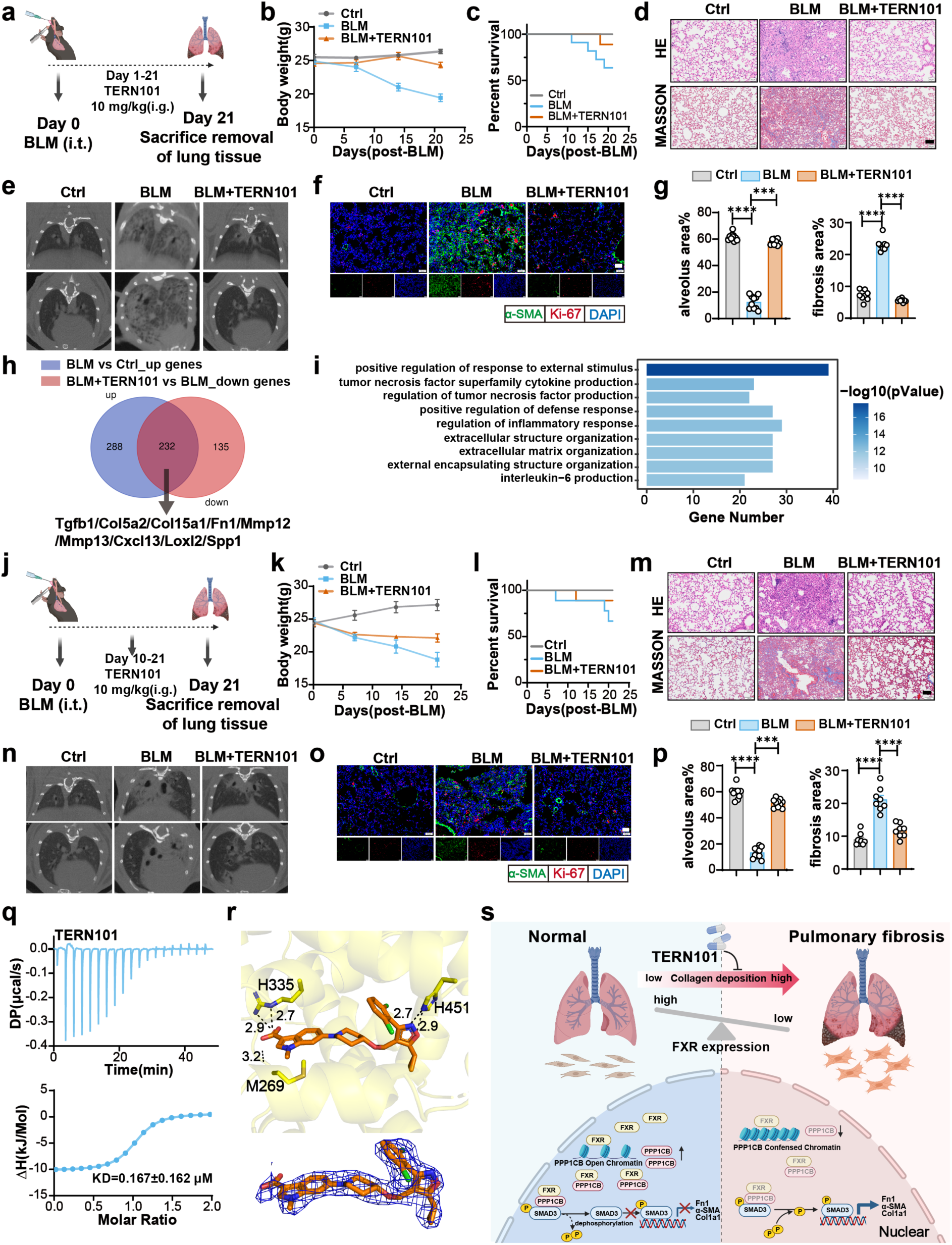
Pharmacological activation of FXR attenuates pulmonary fibrosis progression in mice. (**a**) A Schematic figure of the preventive study. (**b**) Body weight changes in WT mice after BLM challenge with or without TERN101 treatment (n = 5 mice per group). (**c**) Survival curves of WT mice (n = 10 mice per group). (**d, e and g**) Histopathological assessment of pulmonary fibrosis in BLM-challenged mice by H&E staining, Masson’s trichrome (**d,g**), and micro-CT images (**e**). Scale bars: 50 μm (H&E/Masson), 1 mm (micro-CT). (**f**) Representative immunofluorescence images of lung tissue (Red: ki67; Green: α-SMA; Blue: DAPI) (Scar bar: 50 μm). (**h**) Venn diagram. (**i**) Pathway enrichment analysis showing BLM-upregulated pathways inhibited by TERN101. (**j**) Schematic figure of the therapeutic study. (**k-l**) Body weight changes(**k**) and survival curves(**l**) in WT mice after BLM challenge with or without TERN101 treatment (n = 5 mice per group). (**m, n and p**) Histopathological assessment of pulmonary fibrosis in BLM-challenged mice by H&E staining, Masson’s trichrome (**m, p**), and micro-CT images (**n**). Scale bars: 100 μm (H&E/Masson), 1 mm (micro-CT). (**o**) Representative immunofluorescence images of lung tissue (Red: Ki67; Green: α-SMA; Blue: DAPI) (Scar bar: 50 μm). (**q**) Quantitative analysis of ligand-binding thermodynamics for FXR-LBD with TERN101 by isothermal titration calorimetry (ITC). (**r**) Crystal structure of FXR LBD bound to the TERN101 and SRC1-3 peptide (PDB 9WF5). **(s)** The schematic was created in BioRender. Wagner, D. (2025) https://BioRender.com/nmwzx11. The data were assessed by two-tailed Student’s t test. *\*p* < 0.05; *\*\*p* < 0.01; *\*\*\*p* < 0.001; *\*\*\*\*p* < 0.0001; ns, not significant. Data were presented as mean ± SEM.

To elucidate the role of FXR activation in pulmonary fibrosis progression, we established BLM-induced pulmonary fibrosis model in C57BL/6 mice. In a prophylactic intervention setting, the FXR agonist TERN101 was administered daily starting 24 hours after BLM treatment, and lung fibrosis was assessed on day 21 days (**Fig. 6a**). TERN101 significantly mitigated BLM-induced pathophysiology, as evidenced by reduced body weight loss (**Fig. 6b**), improved survival rates (**Fig. 6c**). Histological analysis using H&E and Masson’s staining confirmed attenuation of alveolar structural damage and collagen deposition (**Fig. 6d, g**). which was corroborated by micro-CT imaging (**Fig. 6e**). Immunofluorescence further revealed TERN101 reduced α-SMA and ki-67 expression (**Fig. 6f**). Transcriptomic profiling of lung tissues showed that TERN101 reversed 232 BLM-induced upregulated genes, including key fibrotic regulators such as Tgfb1, Col5a1, Fn1, and Loxl2 (**Fig. 6h, Supplementary Fig. 9a**). KEGG pathway enrichment analysis indicated suppression of profibrotic pathways, including collagen fibril organization, ECM-receptor interaction, and inflammatory signaling (**Fig. 6i, Supplementary Fig. 9b**). Collectively, these findings demonstrate that prophylactic FXR activation by TERN101 effectively prevents pulmonary fibrosis progression.

We next established a therapeutic intervention model in which TERN101 treatment was initiated on day 10 post-BLM exposure (**Fig. 6j**). TERN101-treated mice exhibited attenuated body weight loss (**Fig. 6k**) and improved survival (**Fig. 6l**). Comprehensive pulmonary function tests demonstrated markedly decreased RI and improved FVC, Cchord, and FRC (**Supplementary Fig. 9c**). Histological analysis (**Fig. 6m, p**) and micro-CT imaging (**Fig. 6n**) revealed TERN101-treated marked reductions in structural damage and fibrotic burden, while immunofluorescence confirmed reduced α-SMA and ki-67 expression (**Fig. 6o**).

To validate FXR dependence, we evaluated the efficacy of TERN101 in BLM-treated *Fxr^−/−^* mice (**Supplementary Fig. 9e**). Compared with the BLM-induced model group, TERN101 treatment did not significantly ameliorate the reduction in body weight (**Supplementary Fig. 9f**) and improve survival rate (**Supplementary Fig. 9g**). Furthermore, assessments of pulmonary function (**Supplementary Fig. 9d**), histological examination (H&E and Masson staining) (**Supplementary Fig. 9h, k**), and micro-CT imaging (**Supplementary Fig. 9i**) collectively demonstrated no marked improvement in lung fibrosis and parenchymal damage following TERN101 administration. Immunofluorescence analysis further showed persistent α-SMA and ki-67 expression (**Supplementary Fig. 9j**). Overall, these results confirm that the antifibrotic effects of TERN101 are critically dependent on FXR, reinforcing its role as a critical regulator of pulmonary fibrogenesis.

We subsequently investigated whether other FXR agonists, such as OCA, also exert anti-fibrotic effects. To elucidate the impact of OCA on the progression of pulmonary fibrosis, we established a BLM-induced fibrosis model in C57BL/6 mice. In a preventive intervention regimen, the FXR agonist OCA was administered daily starting 24 hours after BLM exposure, and pulmonary fibrosis was assessed on day 21 (**Supplementary Fig. 11a**). OCA treatment ameliorated BLM-induced body weight loss in mice (**Supplementary Fig. 12b**). Histological analysis using H&E and Masson staining confirmed attenuated alveolar structural damage and collagen deposition (**Supplementary Fig. 13c**). Immunofluorescence further revealed decreased expression of α-SMA and Ki-67 (**Supplementary Fig. 14d**). Collectively, these results demonstrate that different FXR agonists exhibit consistent anti-fibrotic effects, further highlighting the role of FXR as a key regulator in the progression of pulmonary fibrosis.

## Discussion

Pulmonary fibrosis is a devastating progressive lung disease with limited therapeutic options, where increased TGF-β levels prompt fibroblasts to differentiate into myofibroblasts, enhancing matrix production and deposition^44–46^. FXR is a member of the nuclear receptor (NR) superfamily^47^. In this study, we demonstrate that FXR negatively regulates the TGF-β/SMAD signaling pathway by recruiting PPP1CB to mediate SMAD2/3 dephosphorylation. Notably, FXR agonist TERN101 demonstrated therapeutic efficacy in preclinical in vivo models. Collectively, these findings establish FXR as a negative regulator of pulmonary fibrosis and identify FXR as a promising therapeutic target for IPF.

FXR is a bile acid-activated nuclear receptor that regulates multiple physiological and pathological processes, including bile acid homeostasis, glucose and lipid metabolism, and fibrosis^48^. Its role in liver^24^ and kidney fibrosis has been established^24,25^. For example, FXR activation also induces SHP, which inhibits AP-1 activity via JunD binding, reducing collagen expression in hepatic stellate cells^24,49^ In diabetic nephropathy, FXR activation attenuates renal fibrosis by suppressing lipid metabolic dysregulation and profibrotic factors^25^. Of note, FXR may also participate in key pathophysiological processes in the lung, beyond its established roles in the liver and kidney. Nevertheless, existing studies on FXR in fibrotic diseases have focused predominantly on the digestive and urinary systems. Accordingly, the role of FXR in pulmonary fibrosis remains controversial, and its molecular mechanisms are largely unknown. This knowledge gap prompted us to systematically investigate the function of FXR in pulmonary fibrosis. In this study, we found that FXR expression is significantly reduced in lung tissues from IPF patients, and its expression level correlates positively with pulmonary function, suggesting that FXR may act as a protective factor in the regulation of pulmonary fibrosis. Mechanistically, we demonstrate that FXR activation suppresses SMAD2/3 phosphorylation, thereby downregulating the expression of fibrotic genes and inhibiting myofibroblast activation. These findings initially seem at odds with previous observations. Earlier studies reported that certain bile acids (LCA, CDCA) promote IPF through both FXR-dependent and FXR-independent pathways, inducing EMT and activating lung fibroblasts^49^. CDCA has also been implicated in NLRP3 inflammasome activation^50^. However, those studies used natural bile acids, whereas our study employed TERN101—a synthetic, non-bile acid FXR agonist with greater selectivity. By enabling precise FXR activation, TERN101 avoids the receptor promiscuity and concentration-dependent bidirectional effects of natural bile acids, resulting in superior anti-fibrotic efficacy. Nevertheless, it should be acknowledged that the clinical translation of FXR agonists remains challenging. Several FXR agonists, including obeticholic acid and TERN101, have encountered limitations during clinical development due to adverse effects such as pruritus and metabolic disturbances, raising concerns regarding their long-term safety and tolerability. Importantly, recent work has further highlighted the therapeutic potential of FXR by demonstrating that the limitations of current FXR agonists may arise from sustained rather than physiological pulsatile receptor activation. Using the short-acting FXR agonist linafexor, the authors showed that transient, pulsatile FXR activation preserves receptor responsiveness while reducing toxicity and maintaining robust therapeutic efficacy across multiple liver fibrosis models^51^. Collectively, these findings further reinforce FXR as a promising therapeutic target and support the development of next-generation FXR agonists.

Pulmonary fibrosis is a multicellular process involving fibroblasts, type I/II alveolar epithelial cells, and endothelial cells^52^. To define the predominant cell types expressing FXR in fibrotic lungs, we re-analysed published single-cell RNA-sequencing datasets from patients with pulmonary fibrosis. FXR expression was enriched in fibroblasts and mesothelial, implicating FXR plays a crucial role in fibroblast. Fibroblasts are key effector cells that drive fibrotic progression by producing and remodeling the ECM, including collagens, glycoproteins^33,53^. Therefore, elucidating the function of FXR in these cells is of particular importance. In addition, previous studies have implicated FXR in the regulation of pulmonary endothelial cell function. In an LPS-induced acute lung injury model, FXR was shown to reduce endothelial inflammation and vascular permeability and to promote repair^54^. The development and progression of pulmonary fibrosis involve coordinated actions and crosstalk among multiple cell types, we used *Fxr ^−/−^* mice to evaluate the impact of FXR deficiency in BLM-induced pulmonary fibrosis. Compared with BLM-treated WT controls, *Fxr ^−/−^* mice developed more severe lung pathology, supporting an overall protective, anti-fibrotic role for FXR. However, the global knockout model cannot delineate cell type–specific contributions, and it remains unclear whether the phenotype is driven primarily by fibroblasts. This is also one of the limitations of the present study.

Given that FXR lacks intrinsic kinase or phosphatase activity, we hypothesized that FXR might regulate SMAD2/3 phosphorylation by interacting with protein kinases or phosphatases. We further explored the mechanism of FXR function and screened for potential interacting proteins in primary myofibroblasts derived from IPF patients. We focused our investigation on PPP1CB, a key member of the serine/threonine protein phosphatase family that regulates diverse cellular processes including glycogen metabolism, cell cycle progression, and signal transduction^55–57^. Previous studies have demonstrated that PPP1CB mediates the dephosphorylation of STAT3 upon binding to NSD3, thereby suppressing STAT3 nuclear translocation and HK2 transcription, ultimately reducing glycolysis and proliferation^41^. Similarly, in bladder cancer, PPP1CB maintains YBX1 in a hypoactive state by dephosphorylation, attenuating its transcriptional activation of drug-resistance genes^58^. Co-IP and functional assays, we demonstrate for the first time that the DBD of FXR interacts with PPP1CB, leading to suppressed SMAD2/3 phosphorylation and nuclear translocation. However, DBD deletion did not completely abolish the inhibitory effect of FXR overexpression on SMAD-mediated transcriptional activity, suggesting that FXR may suppress SMAD signaling through additional mechanisms beyond PPP1CB binding. Previous studies have shown that FXR activation induces SHP to inhibit AP-1 activity and reduce collagen expression in hepatic stellate cells^24^. FXR overexpression upregulated SHP expression in MRC-5 cells (**Supplementary Fig. 5j**). We therefore speculate that FXR-mediated SHP regulation may also play a role in pulmonary fibrosis, a possibility requiring further investigation. Notably, ATAC-seq analysis revealed reduced chromatin accessibility at the PPP1CB locus in FXR KO MLFs. However, the precise mechanism by which FXR enhances PPP1CB transcriptional activity remains unresolved and warrants further investigation.

In conclusion, our results demonstrated that genetic ablation of FXR exacerbates BLM-induced pulmonary fibrosis by promoting fibroblast hyperactivation and dysregulation of the TGF-β/SMAD signaling pathway. FXR overexpression enhances chromatin accessibility at the PPP1CB locus, facilitating its transcriptional upregulation. This facilitates the formation of a functional FXR-PPP1CB complex, which subsequently suppresses SMAD2/3 phosphorylation and nuclear translocation, thereby inhibiting myofibroblast activation and downstream profibrotic signaling (**Fig. 6s**). Notably, FXR clinical-stage compound TERN101 exhibits significant therapeutic efficacy in a mouse model of pulmonary fibrosis. This study identifies a novel therapeutic target for pulmonary fibrosis and provides new directions for drug development.

## Methods

Our research complies with all relevant ethical regulations.

### Cell Culture

HEK293T, A549 and MRC5 were purchased from Precella. All cell lines were authenticated via STR profiling. HEK293T and A549 cells were cultured in DMEM (Gibco). MRC5 cells were cultured in MEM (Gibco). Primary mouse lung fibroblasts were cultured in DMEM/F12. IPF patient-derived primary myofibroblasts were cultured in DMEM (Gibco). All media were supplemented with 10% FBS (Gibco), 1% penicillin/streptomycin (Gibco). The cells were cultured in 5% CO_2_ at 37 °C. The cells were passaged at 1:3 using 0.25% Trypsin (Gibco) when they reached 80-90% confluence.

### Mice

All mice were maintained in a temperature-controlled facility with 12h light/dark cycle at 23±3°C and 30–70% humidity. All animal procedures were approved by the Institutional Animal Care and Use Committee of Guangzhou Laboratory Animal Center (Protocol No. GZLAB-AUCP-2022-10-A10).

### Generation of *Fxr ^−/−^*mice

*Fxr ^−/−^* mice were generated by Shanghai Model Organisms Center, Inc using CRISPR/Cas9 strategy. Briefly, gRNA1, gRNA2, gRNA3, gRNA4, and Cas9 expression plasmids were designed to delete exon 9 of FXR. Cas9 mRNA and gRNAs were microinjected into fertilized eggs. All mice were compared with non-transgenic or wild-type gender-matched littermates. The primers used for verifying the genotypes of mice are: forward 5’-AGAGCCACCTGGCATTACAC-3’ and reverse 5’-TGGGCAGCCATAGTGTTGAG-3’.

### Patients and Samples Collection

All experiments with human tissue were performed under protocols approved by the Institutional Review Boards at Guangzhou National Laboratory and Ethics Committee of Guangzhou Medical University. Human lung tissue was obtained from patients with IPF (n=16) who underwent lung transplantation. Control lung tissues (n=18) were sourced from the following two groups: (a) histologically normal tissue procured from the non-cancerous regions of lungs following surgical resection for lung cancer, and (b) donor lungs that had been rejected for transplantation for reasons unrelated to IPF. Due to the policy regarding anonymization of data from healthy donors, clinical characteristics are presented for only 10 patients with IPF (Supplementary Table 4). Written informed consent was obtained from each participant before inclusion in the study. All research involving human participants was performed in compliance with all relevant ethical guidelines and was approved by an institutional review board.

### Fibroblast isolation and culture

A piece of fresh lung tissue was weighed, minced into small pieces (1-2 mm^3^). The tissue was digested in DMEM/F12 medium (Gibco) containing 1 mg/mL collagenase I (C9407-sigma) for 2 h in 37℃. The digested tissue pieces and the medium were transferred into the filter (70 μm) and centrifugated at 400g for 5 mins. The DMEM/F12 medium with 10% fetal bovine serum was added for further culture at 37℃ and 5% CO2. IPF-derived myofibroblasts were cultured in DMEM containing 10% FBS (Gibco) and 1% penicillin/streptomycin (Gibco) at 37 °C and tested for mycoplasma regularly. All experiments were carried out using cells at low cell passages (<P4) and were serum-starved during treatment.

### Antibodies

The primary antibodies were used: For Western blot: Fibronectin Polyclonal antibody (1:2000, Proteintech, cat.no.15613-1-AP), anti-COL1A1 (1:1000, Cell Signaling Technology, cat.no. E8F4L), anti-α-SMA (1:1000, Abcam, cat.no. ab7817), anti-phospho-SMAD2 (Ser465/467)/SMAD3 (Ser423/425) (1:1000, Cell Signaling Technology, cat.no. D27F4), anti-Smad2 + Smad3 (1:1000, Abcam, cat.no. ab202445), anti-NR1H4(1:2000, Proteintech, cat.no.25055-1-AP), anti-V5 tag rabbit antibody (1:2000, Proteintech, cat.no.14440-1-AP), anti-V5 tag mouse antibody (1:5000, Abclonal, cat.no. AE017), anti-GAPDH (1:5000, Proteintech cat.no.HRP-60004), anti-PPP1CB (1:1000, Proteintech, cat.no. 10140-2-AP), Histon H3 (1:5000, Proteintech, cat.no. 17168-1-AP). For IP: anti-NR1H4 (Proteintech, cat.no.25055-1-AP), anti-V5 tag rabbit antibody (Proteintech, cat.no.14440-1-AP), anti-V5 tag mouse antibody (Abclonal, cat.no. AE017), anti-PPP1CB (1:1000, Proteintech, cat.no. 10140-2-AP), 2–3 μg of the primary antibody was used for each sample. For IF: anti-α-SMA (1:200, Abcam, cat.no. ab7817), anti-COL1A1 (1:200, Cell Signaling Technology, cat.no. E8F4L), anti-phospho-SMAD2 (Ser465/467)/SMAD3 (Ser423/425) (1:100, Cell Signaling Technology, cat.no. D27F4), anti-NR1H4(1:200, Proteintech, cat.no.25055-1-AP), anti-PPP1CB (1:200, Proteintech, cat.no. 10140-2-AP). For CUT&Tag: anti-Smad2 + Smad3 (Abcam, cat.no. ab202445), anti-NR1H4 (Proteintech, cat.no.25055-1-AP), 2 μg of the primary antibody was used for each sample.

### Cell viability

Low-passage primary mouse fibroblasts were plated in 384-well plates at a density of 6,000 cells per well. The cells were treated with three compounds (OCA, GW4064, and TERN101) at eight serially diluted concentrations starting from 50 μM, using a 1:2 dilution ratio. Following 48 hours of treatment, cell viability was quantified using the CellTiter-Glo (CTG) luminescent cell viability assay, and luminescence was measured on a BioTek Synergy Neo multimode plate reader.

### Drug treatment

For fibrosis induction experiments, cells were serum-starved in DMEM containing 0.5% FBS for 24 h. Subsequently, cells were treated with recombinant human TGF-β (10 ng/ml; Proteintech, cat. no. HZ-1011) for 48 h. Primary mouse lung fibroblasts were cultured under identical conditions and treated with TGF-β for 48 h. Three FXR agonists - Obeticholic acid (OCA; 10 μM) (MACKLIN, cat. no. I843840), GW4064 (10 μM) (Bide Pharmatech, 132493), and TERN101 (10 μM) (Bide Pharmatech, cat. no. BD01071364) were co-administered with TGF-β throughout the 48-h treatment period. For protein phosphatase 1 (PP1) inhibition studies, cells were treated with okadaic acid (15 nM; MCE, cat. no. HY-N6785) for 24 h. OCA, GW4064, TERN101 were dissolved in DMSO, with final DMSO concentrations maintained below 0.1% (v/v) in culture media. Vehicle controls containing equivalent DMSO concentrations were included in all experiments. Okadaic acid was dissolved in ethanol. Vehicle controls containing equivalent ethanol concentrations were included in all experiments. For p-SMAD2/3 detection, cells were treated with TGF-β for 1 hour.

### Construction of Knockdown and Overexpression Cell Lines

For overexpression, the coding sequences of FXR and PPP1CB were cloned into the pLVX-puro or pLVX-Hygro vector. For gene knockdown, shRNAs targeting specific genes (sequences are provided in Supplementary Table 3) were cloned into the pLKO.1-puro vector (Miaoling, Wuhan). Lentiviral particles were generated by co-transfecting HEK293T cells with the respective expression or shRNA plasmids along with the packaging plasmids psPAX2 and pMD2.G using X-tremeGENE 9 transfection reagent. The viral supernatant was collected 48 h post-transfection and filtered through a 0.45-μm membrane. Target cells were infected with the viral particles for 48 h in the presence of 8 μg/mL polybrene. Following infection, cells were selected with medium containing puromycin or hygromycin. The efficiency of gene knockdown or overexpression was validated using qRT-PCR and Western blot analysis.

### Plasmid Construction, Protein Expression and Purification

Wild type human FXR ligand binding domain (LBD) protein, residues 244-472, was expressed from a pET46 plasmid as a 3C-clevable N-terminal 6×His-tag fusion in *Escherichia coli* BL21 (DE3) cells. Proteins were purified using Ni-NTA affinity chromatography, in some cases a second Ni-NTA affinity chromatography step after cleavage of the 6×His-tag by 3C protease (at a ratio of 1 mg 3C protease: 50 mg protein), and finally gel filtration chromatography. Purified protein was concentrated to 10 mg/mL in a buffer consisting of 20 mM Phosphate buffer (pH 7.4, 50 mM KCl + 5 mM tris(2-carboxyethyl) phosphine (TCEP),0.5 mM ethylenediaminetetraacetic acid (EDTA). Purified proteins were verified by SDS-PAGE as >95% pure.

### Crystallization and structure determination

Purified FXR-LBD was purified as described above. FXR-LBD protein was incubated at a 1:3 protein/ligand molar ratio in tris buffer (25 mM Tris-HCl +100 mM NaCl + 0.5 mM EDTA + 3 mM TCEP + 10% glycerol, pH 7.8) overnight, then next day protein was incubated at a 1:3 protein/peptide molar ratio overnight and concentrated to 10 mg/mL. Protein complex crystals were obtained after 5–8 days at 22 °C by sitting-drop vapor diffusion against 50 μL of well solution using 96-well format crystallization plates. The crystallization drops contained 1 μL of protein complex sample mixed with 1 μL of reservoir solution containing 0.1 M Potassium thiocyanate, 30% w/v Polyethylene glycol monomethyl ether 2,000. Crystals were flash-frozen in liquid nitrogen before data collection.

X-ray diffraction data were collected at beamline BL19U1 and BL02U1 at the Shanghai Synchrotron Radiation Facility and processed with the program autoPROC1.0^59^ and Aquarium^60^. The structures were solved by molecular replacement using the program Phaser^61^ integrated in PHENIX^62^ package using the previously published FXR structure (PDB code: 3DCT)^63^ as the search model. The structures were refined using PHENIX with several cycles of manually interactive model rebuilding in Coot^64^. The ligand constraints were generated using Phenix.elbow^65^ and the ligand models were built according to the omit map. The data collection and refinement statistics are summarized in Supplementary Table 1.

### Hydrogen-deuterium exchange mass spectrometry (HDX-MS)

Hydrogen-deuterium exchange (HDX) experiments were analyzed by tandem mass spectrometry (MS/MS) on a Q Exactive HF mass spectrometer (Thermo Fisher). Data were acquired in data-dependent mode with the top eight most abundant ions selected for fragmentation per scan. Peptides were identified with high confidence using pFind.

HDX-MS analysis: FXR-LBD was incubated with or without ligand(OCA, GW4064 and TERN101) at a 1:50 molar ratio (protein: ligand) for 30 min at 4 °C prior to HDX. Then, 4 µL of each sample was diluted into 16 µL of D2O in exchange buffer (50 mM HEPES, pH 7.5, 50 mM NaCl) and incubated for various time points (0, 10, 60, 300, 900 s) at 4 °C. The reaction was quenched with 20 µL of ice-cold 3 M guanidine hydrochloride and 1% trifluoroacetic acid. Quenched samples were immediately injected into the LEAP Pal 3.0 HDX system. Upon injection, samples were passed through an immobilized pepsin column (2mm×2cm) at 120µL/min, and the digested peptides were captured on a C18 PepMap300 trap column (Thermo Fisher) and desalted. Peptides were separated with a 2.1 mm x 5cm C18 separating column (1.9 μm Hypersil Gold, Thermo Fisher) with a linear gradient of 4–40% CH3CN and 0.3% formic acid over 6min. Data were acquired on a Q Exactive HF mass spectrometer (Thermo Fisher) at 65,000 resolution (at m/z 400). All HDX experiments were performed in triplicate. Deuterium incorporation was calculated as the intensity-weighted mean m/z centroid shift of peptide ions. Statistical significance was assessed using an unpaired t-test for each time point within HDX Workbench software^66^. Back-exchange was corrected for by applying a reference value of 70% deuterium recovery and accounting for the 80% deuteration level of the exchange buffer.

Data rendering: Deuterium uptake values from overlapping peptides were mapped to individual amino acid values using a residue-level averaging method. For each residue, deuterium incorporation levels and peptide lengths were compiled from all relevant overlapping peptides. A weighting function favoring shorter peptides was applied, and the weighted values were averaged to yield a single deuterium uptake value per residue. The first two residues of each peptide and all prolines were excluded.

### Time-resolved fluorescence resonance energy transfer assay (TR-FRET)

Time-resolved fluorescence resonance energy transfer (TR-FRET) assays were performed in low-volume black 384-well plates using 22.5 μL final total volume. For the coregulator interaction assay, each well contained 4 nM 6×His-tagged FXR-LBD,1 nM Tb-anti-HIS antibody (Thermo Fisher), and 400 nM FITC-labeled SRC2-2 peptide in a buffer consisting of 20mM potassium phosphate (PH = 8), 50 mM potassium chloride, 5 mM TCEP, and 0.005% Tween 20. Plates were incubated at room temperature for 1 h and read using Synergy Neo plate reader (BioTek). The Tb donor was excited at 340 nm, its emission was monitored at 495 nm, and the acceptor FITC emission was measured at 520 nm. Data were analyzed using GraphPad Prism by calculating the TR-FRET ratio (520 nm/495 nm).

### Isothermal titration calorimetry (ITC)

ITC experiment was carried out on a PEAQ-ITC-Cell (Malvern Panalytical Limitited/UK) using the ITC200 software (v 1.24.2) for instrument control and data acquisition. OCA, GW4064, TERN101 (present in the syringe at 20 μM) was diluted in a buffer consisting of 20 mM Phosphate buffer + 50 mM KCl +5 mM TECP +0.5 mM EDTA, and 0.1% DMSO. FXR-LBD protein (present in the sample cell at 200 μM) as diluted in the same buffer. Rotational stirring set at 5 μcal s^−1^ and 1200 rpm. Data analysis was performed using software packages NITPIC, SEDPHAT, and GUSSI.

### Luciferase reporter assay

HEK293T cells were cultured in DMEM (Gibco) supplemented with 10% fetal bovine serum (FBS) and 1% penicillin and streptomycin. HEK293T cells were seeded in 10-cm^2^ cell culture well (Corning) at 0.5million for transfection using X-tremeGENE 9 DNA Transfection Reagent (Merck) and Opti-MEM reduced serum media (Gibco) with full-length FXR expression plasmid and 3 × FXRE-luciferase. After incubation for 10-18h, cells were transferred to white 384-well cell culture plate at 5000 cells. After 4 h incubation, cells were treated in triplicate with 20 μL of vehicle control (0.4% DMSO in DMEM media), 3-fold serial dilution of ligands for dose response experiments. Following 24 hours of compound treatment, cells were harvested with 20 μL Britelite Plus (PerkinElmer), and luminescence was measured on a BioTek Synergy Neo multimode plate reader. Data were plotted in GraphPad Prism as luminescence vs. ligand concentration and fit to a sigmoidal dose response curve.

### HEK293T CAGA12 luciferase

The construction of the HEK293T-CAGA12 stable cell line is detailed in the section “Construction of Knockdown and Overexpression Cell Lines”. Overexpression or knockdown of FXR and PPP1CB in this cell line was achieved via lentiviral transduction, as detailed in the same section. Cells were seeded in 96-well or 384-well plates and serum-starved for 24 hours. Subsequently, the cells were treated with TGF-β (5 ng/mL, Proteintech, cat. no. HZ-1011) and the test compounds simultaneously for 24 hours. For FXR-overexpressing HEK293T-CAGA12 stable cells, TGF-β was used at a concentration of 2 ng/mL. Following the treatment period, luciferase value was evaluated by dual-luciferase reporter assay (PerkinElmer). Data were plotted in GraphPad Prism as luminescence vs. ligand concentration and fit to a sigmoidal dose response curve.

### Western blot

Lung tissue specimens and fibroblast cultures were homogenized in RIPA lysis buffer (Proteintech, cat. no. PR20035) supplemented with 1% (v/v) protease inhibitor cocktail (Selleck, cat. no. B14001). Protein concentrations were quantified using the BCA Protein Assay Kit (Thermo Fisher Scientific, cat. no. 23225) according to the manufacturer’s protocol. Equal amounts of protein (30-50 μg per lane) were resolved by SDS-PAGE under reducing conditions. Separated proteins were transferred onto methanol-activated PVDF membranes (Immobilon cat. no. ISEQ00010). The membrane was incubated with the primary antibody overnight at 4°C and then incubated with the secondary anti-rabbit (Proteintech, cat. no. SA00001-2) or anti-mouse (Proteintech, cat No. SA00001-1) antibody for 1 h at room temperature. Finally, ECL plus reagent (Proteintech, cat. no. PK10001) was used in the Fluor Chem E imager to detect blotting.

### Nuclear and Cytoplasmic Fractionation

Stable MRC5 cell lines expressing shPPP1CB or FXR overexpression (FXR OE) were generated by lentiviral transduction. Cells were treated with TGF-β (10 ng/mL) in serum-free medium for 1 h, followed by nuclear–cytoplasmic fractionation to assess Smad2/3 phosphorylation levels. Nuclear and cytoplasmic fractions were prepared using a nuclear and cytoplasmic protein extraction kit (Proteintech, cat. no. PK10014) according to the manufacturer’s instructions. Briefly, cells were collected, washed with prechilled PBS, and lysed in nuclear extraction reagent A supplemented protease inhibitor cocktail and phosphatase inhibitors cocktail (Selleck, cat. no. B14001, cat. no. B15001). After homogenization, the lysate was centrifuged at 6500 × g for 5 min at 4°C, and the supernatant was collected as the cytoplasmic fraction. The pellet was washed sequentially with nuclear washing buffer A and B, then resuspended in nuclear extraction reagent B. Following vigorous vortexing and incubation on ice, the sample was centrifuged at 16,000 × g for 10 min at 4°C, and the supernatant was collected as the nuclear fraction. The nuclear fraction was diluted 1:4 with nuclear dilution buffer. Protein concentrations were determined using a BCA protein assay kit (Thermo Scientific. cat. no. A55860).

### Real-time qPCR

RNA from cancer cells was extracted using Trizol reagent (Takara, cat. no. 9108). Complementary DNA was synthesized by using random hexamers and MMLV reverse transcriptase according to the manufacturer’s instructions (Takara). Real-time PCR was carried out using 2× SYBR Green PCR Mix (Accurate Biology, cat. no.AG11701) on an CFX384 Touch Real-Time PCR Detection System (Bio-Rad). The setting procedures were as follows: 95°C for 30 seconds, 95°C for 5 seconds, 60°C for 30 seconds, a total of 40 cycles. Glyceraldehyde phosphate dehydrogenase (GAPDH) was used as an internal control. The relative expression level of target gene was calculated by the 2 −ΔΔCT method. The specific primers used for Real-time PCR are listed in the Supplementary Table 2.

### Transwell assay

Cells chemotaxis was measured using 24-well Nunc (8 µm pore size) transwell inserts (Corning, cat. no. 3422). Cells were seeded (5 × 10^5^ cells/mL) into the upper chamber in FBS-free medium, while the lower chamber contained complete medium with additional 10% FBS as a chemoattractant. Following 48 hours of co-treatment with TGF-β (10 ng/ml) and TERN101(Bide Pharmatech, cat. no. BD01071364), medium was removed and cells in the lower chamber were stained (crystal violet) and imaged using a Nikon microscope (Nikon, Tokyo, Japan) with a ×10 objective.

### Wound healing assay

MLFs were seeded into 12-well plates and cultured until reaching approximately 80% confluence. A linear wound was created by scraping the cell monolayer with a 10-μL sterile pipette tip, and cellular debris was removed by washing with PBS. Cells were then treated with TGF-β (10 ng/mL) and/or TERN101 (10 μM), which were added simultaneously and maintained for 48 h. Wound closure was monitored using the Livecyte automated cell imaging and analysis system, with images captured at the same positions at 0, 24, and 48 h post-scratching. The wound areas were measured using ImageJ software, and cell migration was assessed by calculating the wound closure rate.

### Immunofluorescence

Cells were fixed with 4% (w/v) paraformaldehyde for 15 min at room temperature. Permeabilization was performed using 0.1% (v/v) Triton X-100 (Sigma-Aldrich, cat. no. T8787) in PBS for 15 min at room temperature. After three washes with PBS, cells were incubated with primary antibodies diluted 1:200 in PBS containing 1% (v/v) FBS (Gibco) overnight at 4°C. Following three additional PBS washes, cells were incubated with species-appropriate secondary antibodies: either Goat anti-Rabbit IgG (H+L) Alexa Fluor 555 (1:200; Proteintech, cat. no. RGAR003) or Goat Anti-Mouse IgG (H&L) Alexa Fluor® 488 (1:200; Abcam, cat. no. ab150113) for 1 h at room temperature protected from light. Finally, the cells were washed and mounted using anti-fade mounting media with DAPI. Fluorescent images were captured using Nikon laser confocal microscope. For the co-distribution of two different proteins, we used two antibodies from different species, and a mixture of two secondary antibodies raised in different species (with two different fluorochromes).

### Bleomycin-induced mouse pulmonary fibrosis model

Male C57BL/6 mice (8-10 weeks old; Vital River Laboratories, Beijing, China) were acclimatized for a minimum of 2 weeks in the Guangzhou Laboratory Animal Center under specific pathogen-free conditions. Mice were maintained in a temperature-controlled environment (23-25°C) with 50% relative humidity and a 12 h light/dark cycle, with ad libitum access to food and water. Mice were administered bleomycin hydrochloride (3 mg kg-1 in 50 μL sterile saline; Selleck Chemicals, cat. no. S1214) via intratracheal instillation. Control animals received an equivalent volume of sterile saline (0.9% NaCl). TERN-101 (10 mg/kg; Bide Pharmatech, cat. no. BD01071364) was administered daily by oral gavage in two distinct treatment regimens: (1) Prophylactic model: TERN-101 administration commenced immediately following bleomycin instillation; (2) Therapeutic model: TERN-101 treatment was initiated on day 10 post-bleomycin administration. Vehicle control groups received equivalent volumes of 0.5% (w/v) methylcellulose solution. On day 21, mice were euthanized followed by collection of blood and lung tissue.

### Measurement of lung function in mice

Following completion of the treatment regimen, mice were anesthetized via intraperitoneal injection of Tribromoethanol. Tracheal intubation connected to the Buxco FinePointe Resistance and Compliance system. The system was calibrated according to the manufacturer’s protocol prior to each measurement session. At least three technically valid measurements were obtained for each animal.

### Histologic assessment of pulmonary fibrosis

The lungs were removed at 21 days after BLM or administration. The lungs were harvested, fixed in 4% paraformaldehyde solution in PBS, and embedded in paraffin. The sections were stained hematoxylin eosin staining and Masson trichrome. Whole-slide digital images were acquired at 20× magnification using a slide scanner (Olympus SLIDEVIEW VS200).

### Immunohistochemistry

Lung tissues were fixed in 4% paraformaldehyde for 48 h at room temperature, embedded in paraffin, and cut into 5-µm-thick sections. For immunohistochemical analysis, sections were incubated overnight at 4°C with an anti-FXR antibody (Proteintech, China). This was followed by incubation with a secondary antibody (Servicebio, China) and HRP-conjugated streptavidin. Signal development was performed using a DAB substrate kit (Servicebio, China), and nuclei were counterstained with hematoxylin (Servicebio, China). After dehydration, the slides were mounted with resinous medium. Images were captured using an Olympus SLIDEVIEW VS200 microscope.

### Multiplex immunofluorescence

Multiplex immunofluorescence staining for FXR, α-SMA, and PDGFRα was performed on frozen lung tissue sections. Following methanol fixation, antigen retrieval was performed in Tris-EDTA buffer (pH 8.0) for 15 min. Endogenous peroxidase was blocked with 3% hydrogen peroxide, and sections were blocked with 3% BSA or 10% normal goat serum. Primary antibodies against FXR (1:200, Proteintech, 25055-1-AP), α-SMA (1:200, Abcam, ab7817), and PDGFRα (1:200, Abcam, ab203491) were applied sequentially with overnight incubation at 4°C, followed by HRP-conjugated secondary antibody (Servicebio) and tyramide signal amplification (iF555, iF647, or iF488 Tyramide, Servicebio). Between staining rounds, antibodies were stripped using an antibody stripping solution at 37°C for 30 min. Nuclei were counterstained with DAPI, and images were captured using an Olympus SLIDEVIEW VS200 microscope.

### RNA sequencing

MLF were treated with a combination of TGF-β (10 ng/mL) and TERN101 (10 μM) for 48 hours. After 21 days of TERN101 treatment in the BLM-induced pulmonary fibrosis mouse model, lung tissues were collected from both model control and TERN101-treated groups for RNA seq. Total RNA was extracted according to the manufacturer’s instructions and submitted to Berry Genomics for sequencing on the Illumina NovaSeq 6000 X Plus platform (Berry Genomics), with approximately 6 GB of sequencing data per sample. Differentially expressed genes were identified using DESeq with fold change >2 and an adjusted p value (FDR)<0.01. FastQC (v 0.11.9) was used for sequencing quality assessment. The raw reads were trimmed using Trim-galore (v 0.6.7) to remove adapters. We mapped the clean reads to the mouse reference genome (mm39) using Hisat2 (v 2.2.1). The number of reads mapped to each gene was counted using the featureCounts program of the Subread (v 2.0.1). We performed differentially expressed genes (DEGs) analysis using the DESeq2 (v 1.34.0). Then, we performed Gene Ontology (GO) enrichment analysis and KEGG pathway analysis to DEGs using clusterProfiler (v 4.6.0). The normalized counts were also variance-stabilizing transformed and scaled to plot the heatmap.

### Single cell RNA sequencing

Single-cell RNA sequencing datasets of lung tissues from healthy donors and patients with IPF were obtained from a previously published study^67,68^. After 21 days of BLM treatment, fresh lung tissues were collected from WT and FXR-knockout mice and subjected to scRNA-seq with two biological replicates per group. Two independent scRNA-seq experiments were performed. Whole-lung single-cell suspensions (all cell populations) were directly processed for scRNA-seq. Libraries were sequenced on the DNBSEQ-T7 platform at an average sequencing depth of ∼3,704×.

Raw sequencing data were processed into a gene–cell expression matrix using dnbc4 software following the recommended workflow. Downstream analyses, including normalization, dimensionality reduction, clustering, and cell-type annotation, were performed using Seurat (R), assisted by ACT (http://xteam.xbio.top/ACT/). Transcription factor (TF) regulon activities were inferred using pySCENIC (Python) with default parameters. The inferred regulon activity matrix was imported into the Seurat object as an additional assay for subsequent analyses. Data visualization was primarily performed using the scplotter and scCustomize R packages.

### CUT&Tag sequencing

We performed the CUT&Tag sequencing using the CUT&Tag Assay Kit (Novoprotein). MLF were treated with a combination of TGF-β (10 ng/mL) and TERN101 (10 μM) for 48 hours, followed by overnight incubation with p-SMAD2/3 antibody (abcam) at 4°C. The resulting DNA was amplified to generate sequencing libraries, which were subjected to paired-end 150-bp sequencing on the DNBSEQ-T7 platform with a sequencing depth of approximately 3.70×. We performed quality control using FastQC on all samples. The raw reads were trimmed using Trim-galore to remove adapters. The clean reads were mapped to mouse reference genome (mm39) using Bowtie2 with the “--end-to-end --very-sensitive -I 10 -X 700 --no-mixed” option. Low-quality mapped reads and those mapped to mitochondrial reads were filtered using Samtools. The duplicated reads were removed using Sambamba. The Counts Per Million mapped reads (CPM) normalization method was used to transform alignment bam files into read coverage files in bigWig format on deepTools. Peak calling was performed using MACS2. Differentially accessible peaks were identified using DiffBind. Motif enrichment analysis was performed using HOMER. Further analysis included peak visualization and genomic annotation using IGV (v2.16.2).

### ATAC sequencing

We performed the ATAC sequencing using the ATAC Assay Kit (Vazyme). The resulting DNA was amplified to generate sequencing libraries, which were subjected to paired-end 150-bp sequencing on the DNBSEQ-T7 platform with a sequencing depth of approximately 3.70×. We performed quality control using FastQC on all samples. The raw reads were trimmed using Trim-galore to remove adapters. The clean reads were mapped to mouse reference genome (mm39) using Bowtie2 (v 2.4.5) with the “--very-sensitive -X 1000” option. Low-quality mapped reads and those mapped to mitochondrial reads were filtered using Samtools (v 1.15.1). The duplicated reads were removed using Sambamba (v 0.8.2). The Counts Per Million mapped reads (CPM) normalization method was used to transform alignment bam files into read coverage files in bigWig format on deepTools (v 3.5.1). Peak calling was performed using MACS2 (v 2.2.7.1) with the parameter “--SPMR --nomodel --shift -100 --extsize 200”. Differentially accessible peaks were identified using DiffBind (v 3.8.0). Motif enrichment analysis was performed using HOMER (v 4.11). Further analysis included peak visualization and genomic annotation using IGV (v2.16.2).

### Co-Immunoprecipitation (Co-IP)

Cells were incubated in IP lysis buffer (25mM Tris, 150mM NaCl, 1mM EDTA,1%NP40, 5%Glycerol, PH 7.4). Cell lysates were incubated with antibody overnight at 4℃. Rabbit IgG (Proteintech) was used as a negative control. The next day, protein A/G agarose beads (20422, Thermo fisher) were added to the lysates and incubated for 6h at 4℃. Then the agarose beads were harvested and washed 3 times with PBS, followed by boiling for 10 min at 100 °C. Samples were analyzed by Western blotting to confirm the interaction. The protein samples obtained from Protein A/G bead-based immunoprecipitation were processed through tryptic digestion and desalting procedures before being analyzed by mass spectrometry.

### Statistical Analysis

All statistical analyses were conducted using GraphPad Prism software. The data are represented as mean ± SEM calculated using GraphPad. A two-tailed Student’s t test was utilized to compare two groups, while a one-way ANOVA followed by Tukey’s post hoc test was employed for comparisons involving three or more groups. A *p*-value of less than 0.05 was considered statistically significant.

## Data availability

Crystal structures generated during the current study are available in the Protein Data Bank (PDB) under accession codes 9WF5. The RNA sequencing (RNA-Seq) datasets have been deposited in the Genome Sequence Archive (Genomics, Proteomics & Bioinformatics 2021) in National Genomics Data Center (Nucleic Acids Res 2022), China National Center for Bioinformation / Beijing Institute of Genomics, Chinese Academy of Sciences under accession code CRA030022 that are publicly accessible at https://ngdc.cncb.ac.cn/gsa. Source data are provided with this paper.

## Acknowledgements

We thank Prof. Douglas J. Kojetin, Vanderbilt University, for critical reading and discussions. We are grateful for support from Shanghai Synchrotron Radiation Facility (SSRF) beamlines (BL19U1 and BL02U1) and Guangzhou Laboratory Core Facility and Animal Center. This study was supported by grants from, the startup and Major Program of Guangzhou National Laboratory (GZNL2023A02012 to J.S.; GZNL2025C02004 to J.S.), the National Natural Science Foundation of China (82170473 to J.S.) and the Guangdong Natural Science Foundation (2021QN020451 to J.S.).

## Author contributions

W.L. performed the cellular and biochemical assays. X.L., Y.L. and J.S. performed crystallography. W.L. and X.Y. performed the animal models. W.L. and X.L. performed cloning and purified proteins. W.L. and Q.W. performed mass spectrometry analysis. W.L., M.L., W.H., and W.K. Z performed transcriptome, CUT&Tag and ATAC-seq analysis. X.Q., M.T. and Y.L. performed structural modeling. W.L. and J.Z. performed HDX-MS. W.L. and J.S. designed the experiments, interpreted data, and wrote the manuscript with input from all authors.

