## Supplementary Information for "FXR-mediated recruitment of PPP1CB suppresses SMAD2/3 phosphorylation to mitigate pulmonary fibrosis"

W. Li *et al.*

### Table of Contents

| | Initial page |
| --- | --- |
| **S1: Supplementary Figures** | **3** |
| **S2: Supplementary Tables** | **18** |

#### S1: Supplementary Figures

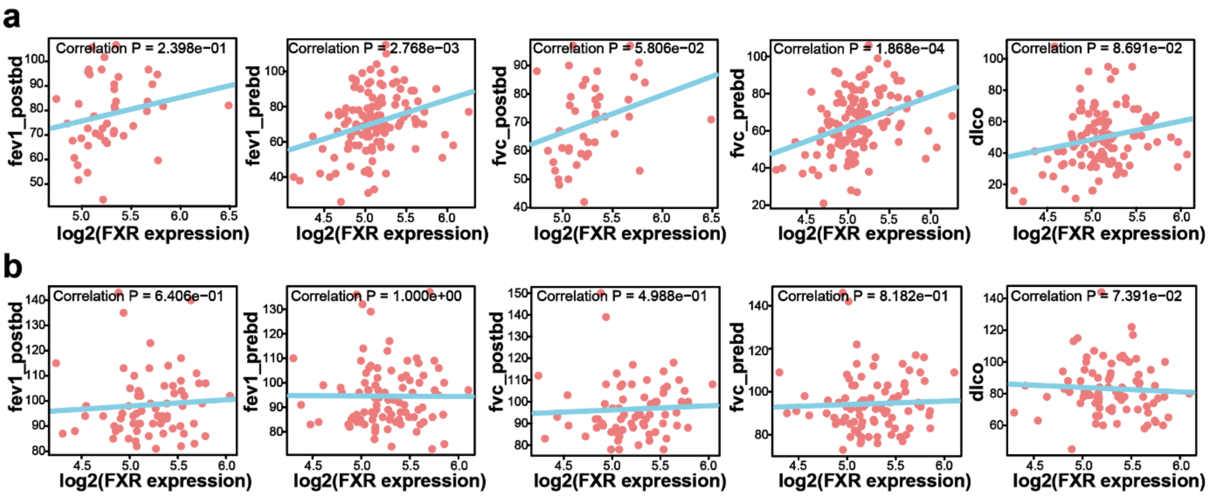

**Supplementary Fig. 1. Decreased expression of FXR is correlated with disease severity of pulmonary fibrosis.** (**a-b**) Correlation analysis between lung function and FXR mRNA levels in lung tissues from patients with IPF(**a**) and healthy controls(**b**). (GEO: GSE47460).

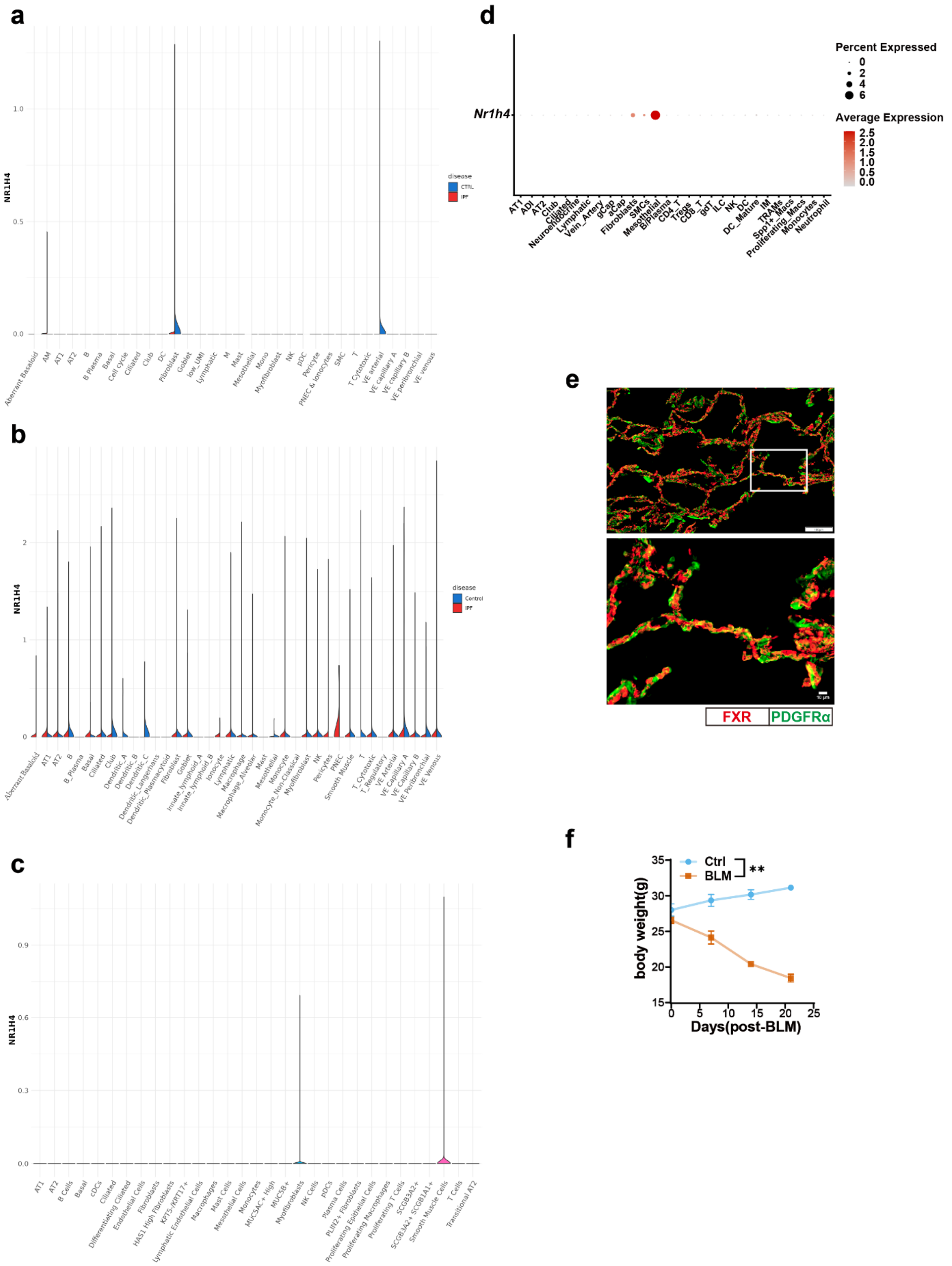

**Supplementary Fig. 2. Decreased expression of FXR is correlated with disease severity of pulmonary fibrosis. (a-b)** FXR expression in lung cell subpopulations from healthy donors and patients with IPF. **(c)** Cell-type–resolved expression of FXR in human IPF lungs based on published scRNA-seq data. (**d**) Expression of FXR across cell populations in BLM-induced fibrotic lung tissue. (**e**) Immunofluorescence showing colocalization of FXR (red) and PDGFR α (green), with scale bars of 100 μm (top) and 10 μm (bottom). (**f**) Body weight changes at day 21 post BLM instillation.

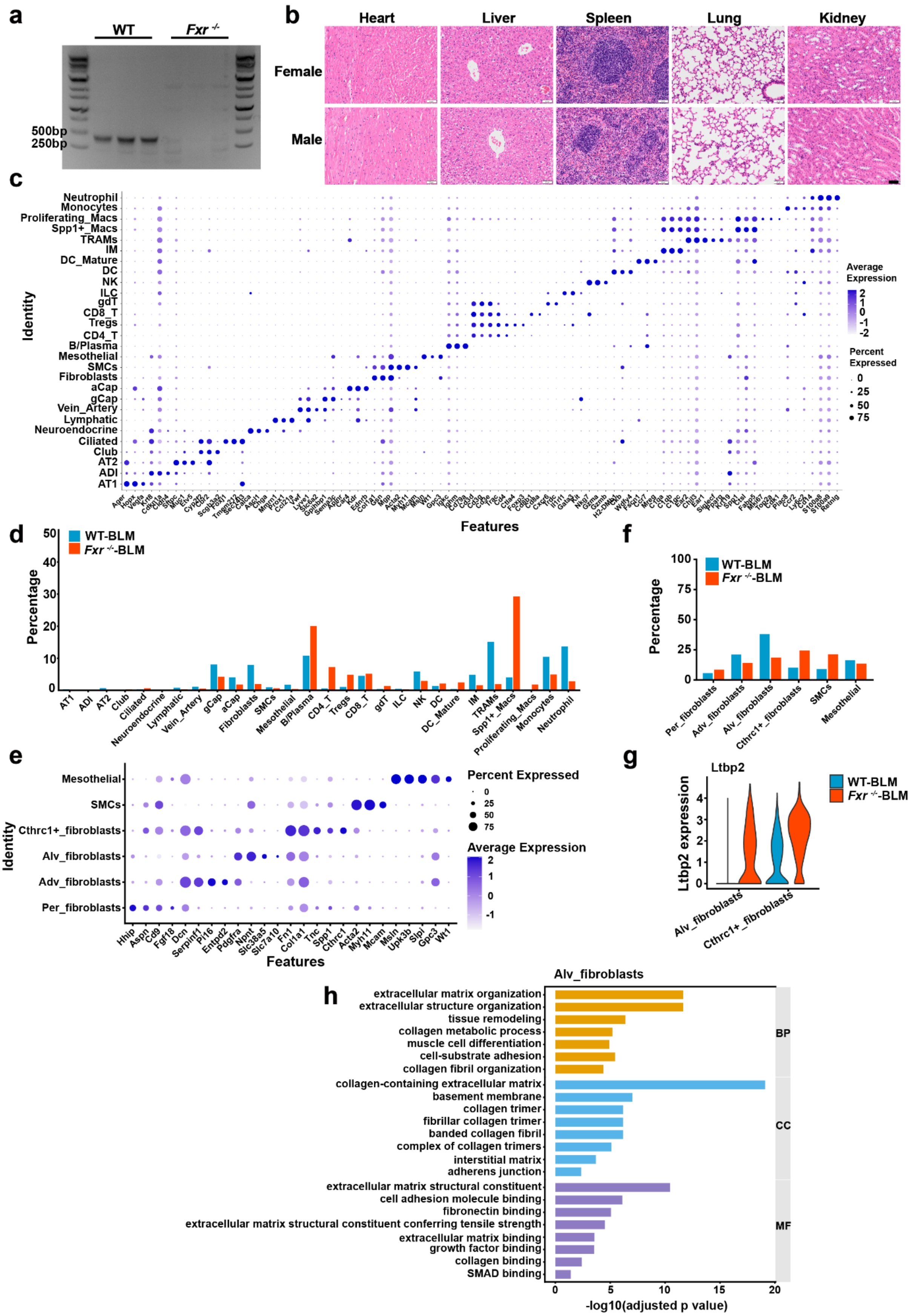

a
WT
Fxr -/-
500bp
250bp
b
Heart
Liver
Spleen
Lung
Kidney
Female
Male
c
Neutrophil
Monocytes
Proliferating_Macs
Spp1+_Macs
TRAMs
IM
DC_Mature
DC
NK
ILC
gdT
CD8_T
Tregs
CD4_T
B/Plasma
Mesothelial
SMCs
Fibroblasts
aCap
gCap
Vein_Artery
Lymphatic
Neuroendocrine
Ciliated
Club
AT2
ADI
AT1
Identity
Features
Average Expression
Percent Expressed
d
WT-BLM
Fxr -/--BLM
Percentage
AT1
ADI
AT2
Club
Ciliated
Neuroendocrine
Lymphatic
Vein_Artery
gCap
aCap
Fibroblasts
SMCs
Mesothelial
B/Plasma
CD4_T
Tregs
CD8_T
gdT
ILC
NK
DC
DC_Mature
IM
TRAMs
Spp1+_Macs
Proliferating_Macs
Monocytes
Neutrophil
e
Mesothelial
SMCs
Cthrc1+_fibroblasts
Alv_fibroblasts
Adv_fibroblasts
Per_fibroblasts
Identity
Hhip
Aspn
Cd9
Fgf18
Dcn
Serpinf1
Pi16
Entpd2
Pdgfra
Npnt
Slc38a5
Slc7a10
Fn1
Col1a1
Tnc
Spp1
Cthrc1
Acta2
Myh11
Mcam
Msln
Upk3b
Slpi
Gpc3
Wt1
Features
Percent Expressed
Average Expression
f
WT-BLM
Fxr -/--BLM
Percentage
Per_fibroblasts
Adv_fibroblasts
Alv_fibroblasts
Cthrc1+_fibroblasts
SMCs
Mesothelial
g
Ltbp2
WT-BLM
Fxr -/--BLM
Ltbp2 expression
Alv_fibroblasts
Cthrc1+_fibroblasts
h
Alv_fibroblasts
extracellular matrix organization
extracellular structure organization
tissue remodeling
collagen metabolic process
muscle cell differentiation
cell-substrate adhesion
collagen fibril organization
collagen-containing extracellular matrix
basement membrane
collagen trimer
fibrillar collagen trimer
banded collagen fibril
complex of collagen trimers
interstitial matrix
adherens junction
extracellular matrix structural constituent
cell adhesion molecule binding
fibronectin binding
extracellular matrix structural constituent conferring tensile strength
extracellular matrix binding
growth factor binding
collagen binding
SMAD binding
BP
CC
MF
-log10(adjusted p value)

**Supplementary Fig. 3. FXR deficiency aggravates pulmonary fibrogenesis in a BLM-challenged mice model.** (**a**) Genotyping results of *Fxr* $^{-/-}$ mice. (**b**) Representative HE staining of mouse organ sections (Scalebar:50 μm). (**c**) Dot plot showing representative markers for each cell cluster. (**d**) Proportions of each cell type in each group.(**e**) Dot plots of fibroblast subtype markers. (**f**) Proportions of fibroblast subtypes across two groups. (**g**) The expression levels of Ltbp2 between the two groups. (**h**) Pathway enrichment analysis in Alv-fibroblasts.

*$p < 0.05$; **$p < 0.01$; ***$p < 0.001$; ****$p < 0.0001$; ns, not significant. Data were presented as mean ± SEM.

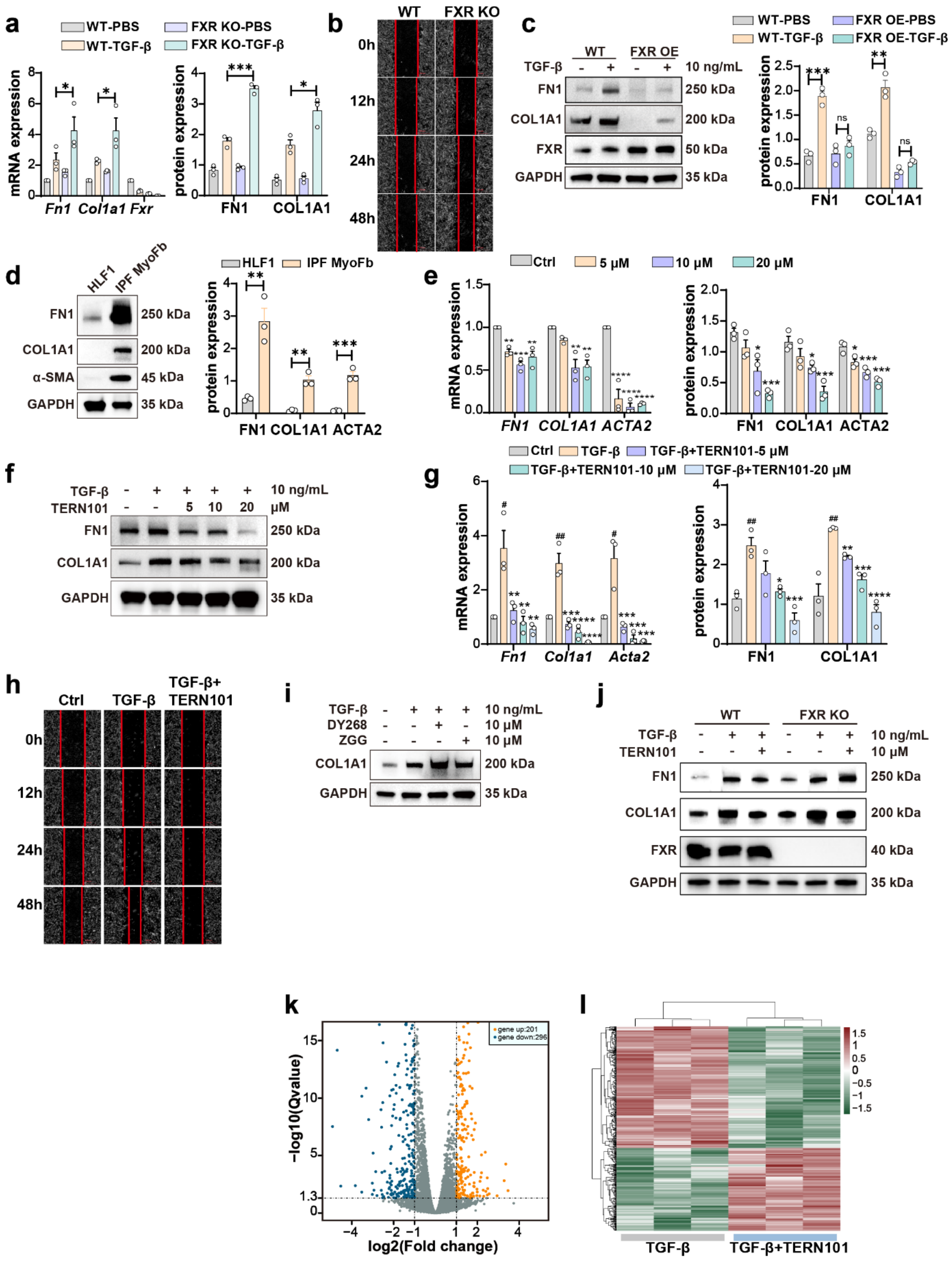

**Supplementary Fig. 4. Pharmacological FXR agonism attenuates myofibroblast activation.** (**a**) Expression level of fibrotic genes in WT and FXR KO MLFs following 48-hour TGF-β stimulation (10 ng/mL) (n=3). (**b**) Wound healing assay showing the migration of WT and FXR KO MLFs. (**c**) Expression level of fibrotic genes in WT and FXR OE MLFs following 48-hour TGF-β stimulation (10 ng/mL) (n=3). (**d**) Expression level of fibrotic genes in HLF1 and IPF MyoFb (n=3). (**e**)

Statistical analysis of fibrotic gene expression in IPF MyoFb treated with TERN101. (**f-g**) The expression of fibrotic genes in MLFs treated with TERN101. (**h**) Wound healing assay showing the migration in MLFs treated with TERN101. (**i**) Western blot analysis of COL1A1 protein level following DY268 (10 μM) and ZGG (10 μM) treatment. . (**j**) Western blot analysis of fibrotic genes protein levels in WT and FXR KO MLFs treated with TGF-β and TERN101. (**k**) A volcano plot showing the differential gene expression between TGF-β and TGF-β+TERN101 in MLFs. (**l**) Ranked plot of TGF-β-responsive genes reversed by TERN101 treatment in MLFs. The data were assessed by two-tailed Student's t test and one-way ANOVA. *$p < 0.05$; **$p < 0.01$; ***$p < 0.001$; ****$p < 0.0001$; ns, not significant. Data were presented as mean ± SEM.

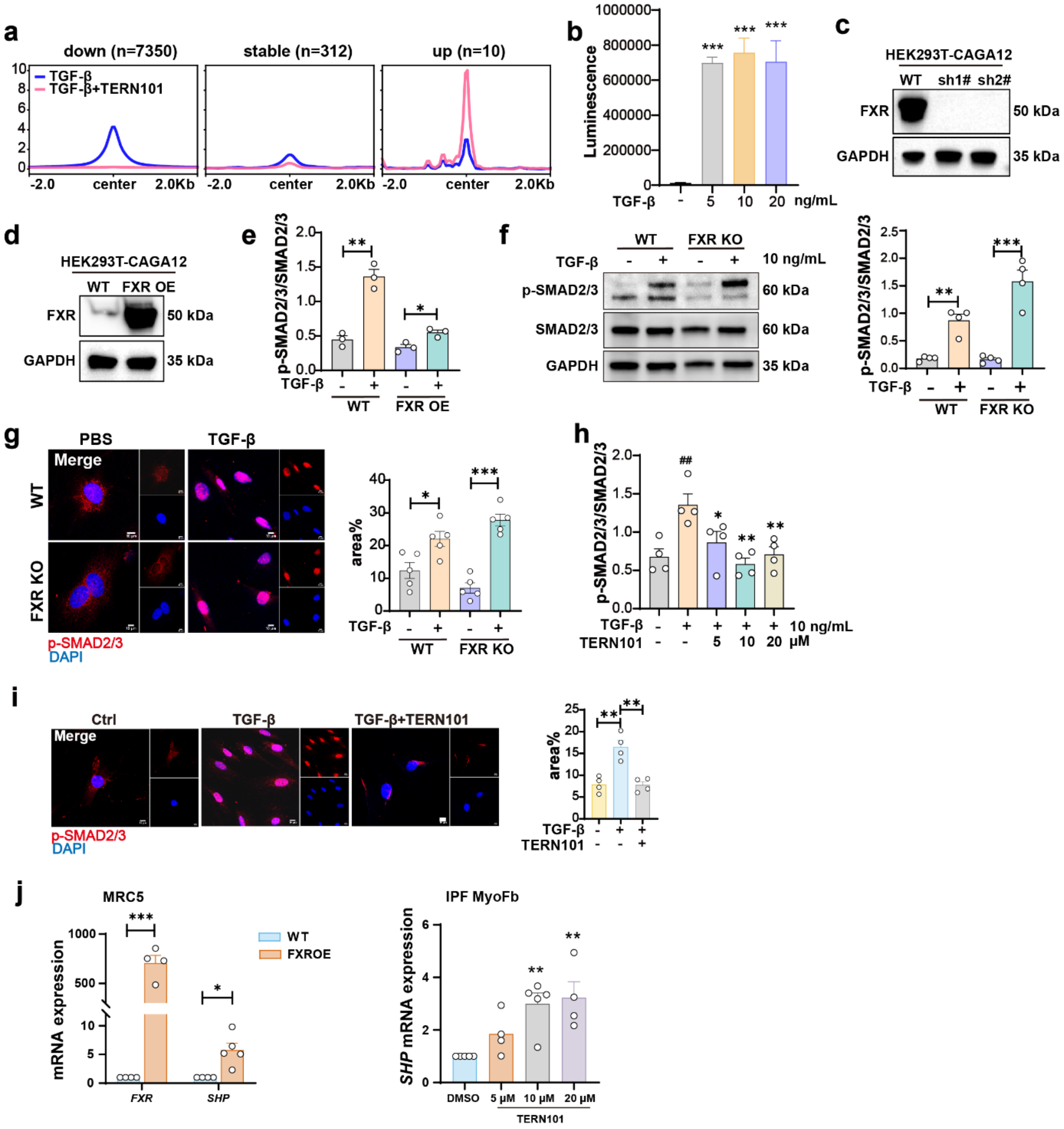

**Supplementary Fig. 5. FXR suppresses SMAD2/3 phosphorylation and nuclear translocation.** (**a**) Average p-SMAD2/3 signal corresponding to (**Fig. 3j**). (**b**) Luciferase reporter assays in HEK293T cells stably expressing the SMAD-responsive CAGA12-luciferase after treatment with TGF-β (n=3). (**c-d**) Western blot analysis of FXR protein expression in HEK293T-CAGA12 cells with FXR knockdown (**c**) and overexpression (**d**). (**e**) Quantitative Western blot analysis of p-SMAD2/3 protein expression in FXR OE MLFs with or without TGF-β (10 ng/mL, 24 h) stimulation. (n=3). (**f**) Western blot analysis of p-SMAD2/3 protein expression in FXR KO MLFs with or without TGF-β (10 ng/ml, 1h) stimulation. (n=4). (**g**) Representative immunofluorescence images of p-SMAD2/3 nuclear translocation in WT and FXR KO MLFs treated with or without TGF-β (10 ng/mL, 1 h). Scale bar: 10 μm. (**h**) Quantitative Western blot analysis of p-SMAD2/3 protein expression in MLFs treated with or without TERN101 (10 μM, 24 h) and TGF-β (10 ng/mL, 1 h) stimulation (n=4). (**i**) Representative immunofluorescence images and quantitative analysis of p-SMAD2/3 nuclear translocation in MLFs treated with or without TGF-β (10 ng/ml, 1h) and TERN101 (10 μM, 24 h) for 24 h. Scale bar: 10 μm. (n=4). (**j**) Effects of FXR overexpression and FXR agonist TERN101 on FXR target gene expression. The data were assessed by two-tailed Student's t test and one-way ANOVA. *$p < 0.05$; **$p < 0.01$; ***$p < 0.001$; ****$p < 0.0001$; ns, not significant. Data were presented as mean ± SEM.

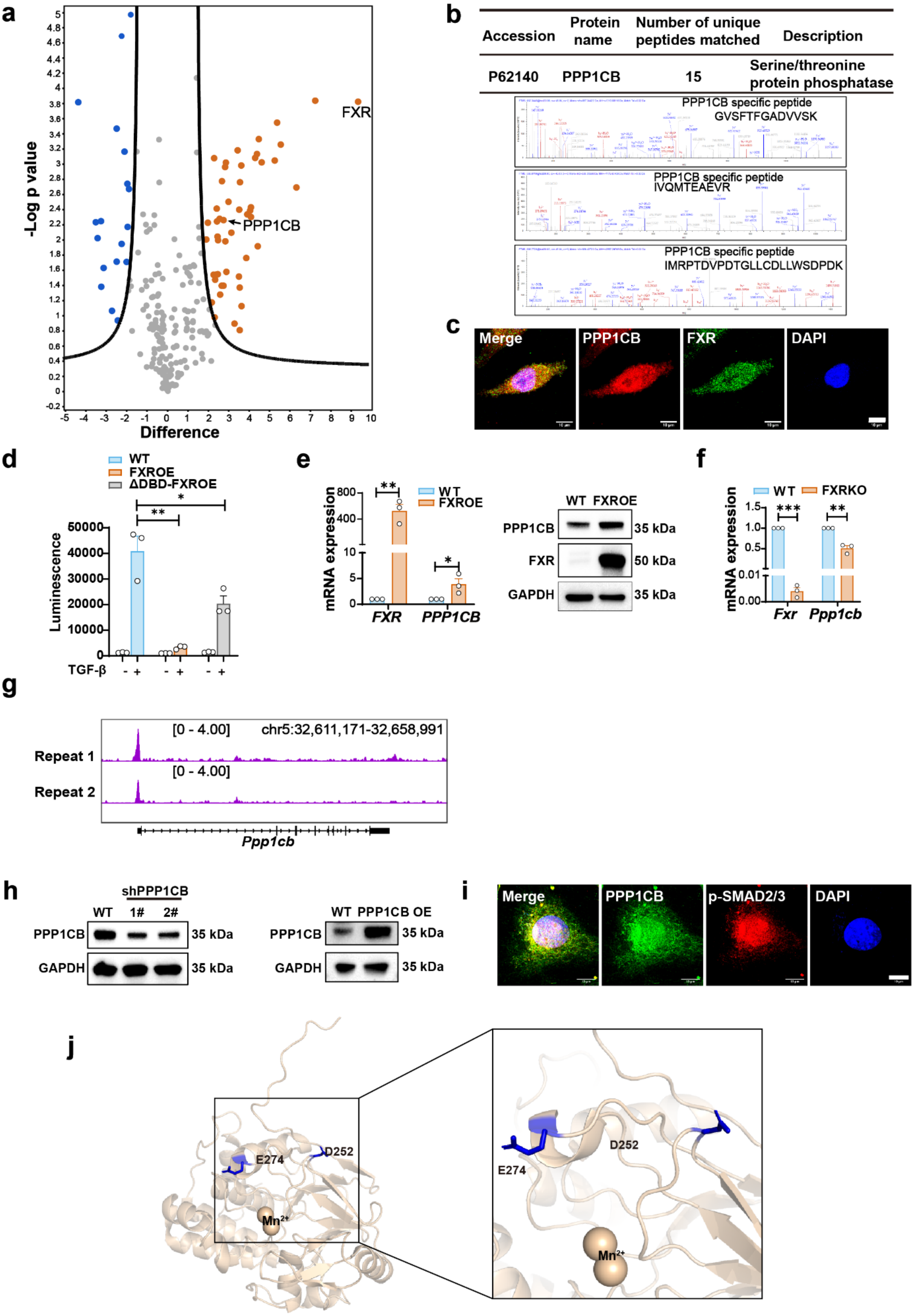

a
-Log p value
Difference
FXR
PPP1CB
b
Accession
Protein name
Number of unique peptides matched
Description
P62140
PPP1CB
15
Serine/threonine protein phosphatase
PPP1CB specific peptide GVSFTFGADVVSK
PPP1CB specific peptide IVQMTEAEVR
PPP1CB specific peptide IMRPTDVPDTGLLCDLLWSDPDK
c
Merge
PPP1CB
FXR
DAPI
d
WT
FXROE
ΔDBD-FXROE
Luminescence
TGF-β
e
mRNA expression
WT
FXROE
FXR
PPP1CB
WT FXROE
PPP1CB
FXR
GAPDH
35 kDa
50 kDa
35 kDa
f
WT
FXRKO
mRNA expression
Fxr
Ppp1cb
g
Repeat 1
Repeat 2
[0 - 4.00]
chr5:32,611,171-32,658,991
[0 - 4.00]
Ppp1cb
h
shPPP1CB
WT 1# 2#
PPP1CB
GAPDH
35 kDa
35 kDa
WT PPP1CB OE
PPP1CB
GAPDH
35 kDa
35 kDa
i
Merge
PPP1CB
p-SMAD2/3
DAPI
j
E274
D252
Mn2+
D252
E274
Mn2+

**Supplementary Fig. 6. FXR-mediated recruitment of PPP1CB inhibits SMAD2/3 phosphorylation.** (**a**) A volcano plot of differential proteins shown in IPMS in WT and FXROE IPF MyoFbs. There were 18 downregulated proteins (blue) and 50 upregulated proteins (orange). (**b**) The PPP1CB specific peptide in the IP product of FXR was discovered through IP/MS analysis. (**c**) Immunofluorescence localization of PPP1CB and FXR in A549 cells. Scale bar: 10 μm. (**d**) Luciferase activity was measured in HEK293T-CAGA12 stable cells overexpressing FXR or FXR-ΔDBD after stimulation with TGF-β (2 ng/mL) for 24 h. (**e**) Expression of PPP1CB in A549 cells with FXR overexpression. (**f**) Expression of PPP1CB in MLFs with FXR knockout. (**g**) IGV analysis of CUT&Tag-seq signals in the gene locus of *Ppp1cb* in MLFs. (**h**) Expression of PPP1CB in HEK293T-CAGA12 cell line with overexpression and knockdown of PPP1CB. (**i**) Immunofluorescence localization of PPP1CB and p-SMAD2/3 in MLFs. Scale bar: 10 μm. (**j**) Overall structure of PPP1CB shown in cartoon representation. The data were assessed by two-tailed Student's t test. *$p < 0.05$; **$p < 0.01$; ***$p < 0.001$; ****$p < 0.0001$; ns, not significant. Data were presented as mean ± SEM.

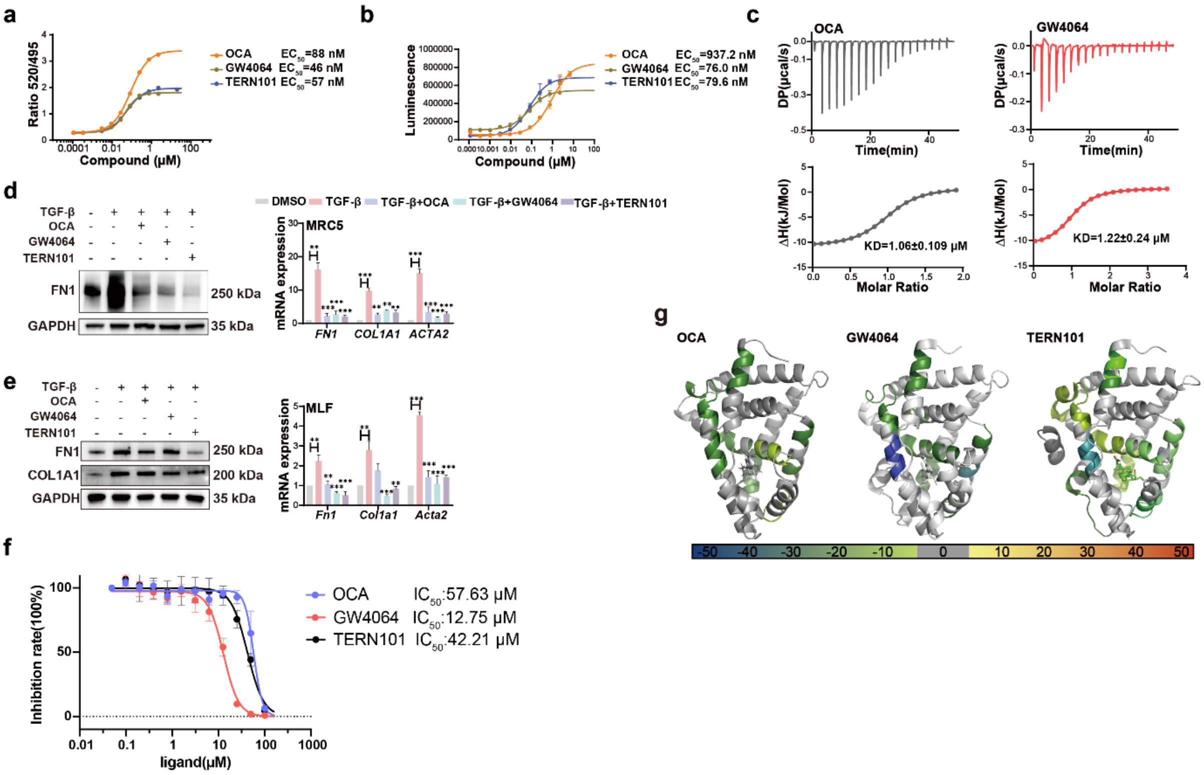

**Supplementary Fig. 7. Pharmacological FXR agonism attenuates myofibroblast activation.** (**a**) TR-FRET assay measuring SRC2-2 peptide recruitment to FXR in the presence of compounds. Data represent FRET signal (520 nm/495 nm ratio) normalized to vehicle control (n=3). (**b**) Luciferase activity in HEK293T cells transfected with FXR-FL and pGL4-FXRE-Luc reporter and treated with ligands for 24 h (n=3). (**c**) Quantitative analysis of ligand-binding thermodynamics for FXR's ligand-binding domain (FXR-LBD) with OCA and GW4064 by isothermal titration calorimetry (ITC). (**d-e**) The effect of FXR agonists on the levels of FN1, COL1A1 and α-SMA proteins and mRNA in both MRC5(**d**) and MLFs(**e**) following TGF-β stimulation (10 ng/ml, 48 h) . (**f**) CTG cytotoxicity assay for quantitative MLFs cell survival analysis (n=5). (**g**) Peptides exhibiting significant changes in hydrogen-deuterium exchange (HDX) upon binding to OCA, GW4064, or TERN101 were mapped onto the FXR crystal structure. The data were assessed by two-tailed Student's t test and one-way ANOVA. *$p < 0.05$; **$p < 0.01$; ***$p < 0.001$; ****$p < 0.0001$; ns, not significant. Data were presented as mean ± SEM.

#### FXR LBD ± TERN101

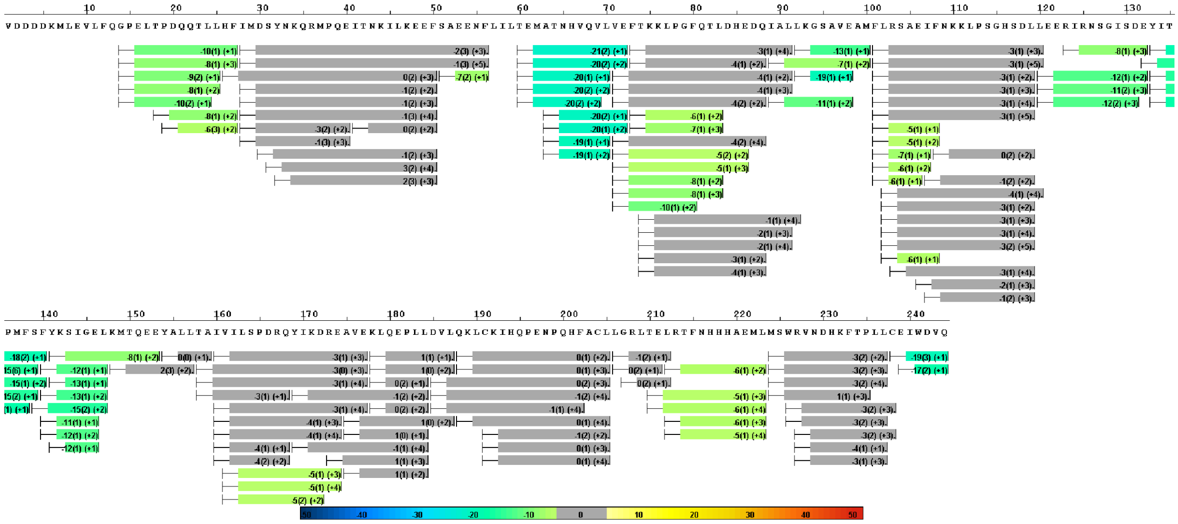

#### FXR LBD ± OCA

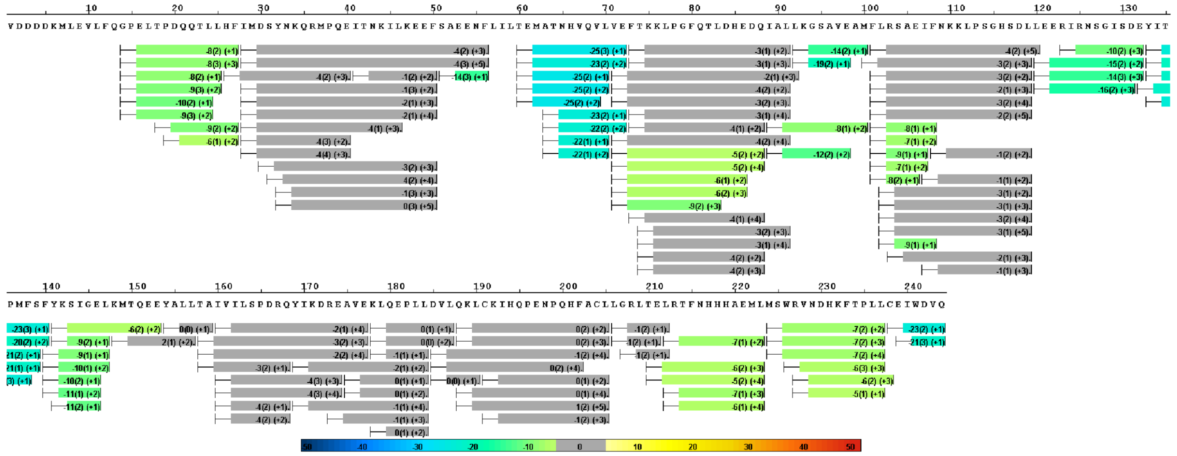

#### FXR LBD ± GW4064

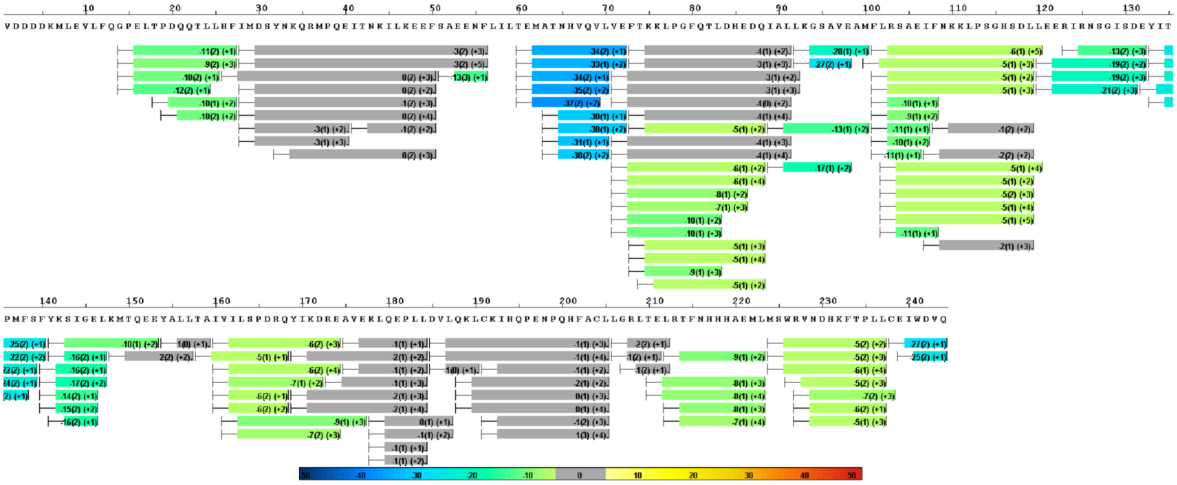

**Supplementary Fig. 8. HDX-MS mapped regions protected by ligand (TERN101、OCA and GW4064) to the peptides in the C-terminal domain of His-tag FXR-LBD (244-472).** Residues with a significantly lower rate of hydrogen–deuterium exchange after TERN101 binding compared to the apo form represented in light green (from -20 to 0 percent of uptake ratio). Residues with a significantly lower rate of hydrogen–deuterium exchange after OCA binding compared to the apo form represented in deep green (from -30 to 0 percent of uptake ratio). Residues with a significantly lower rate of hydrogen–deuterium exchange after GW4064 binding compared to the apo form represented in blue (from -40 to 0 percent of uptake ratio).

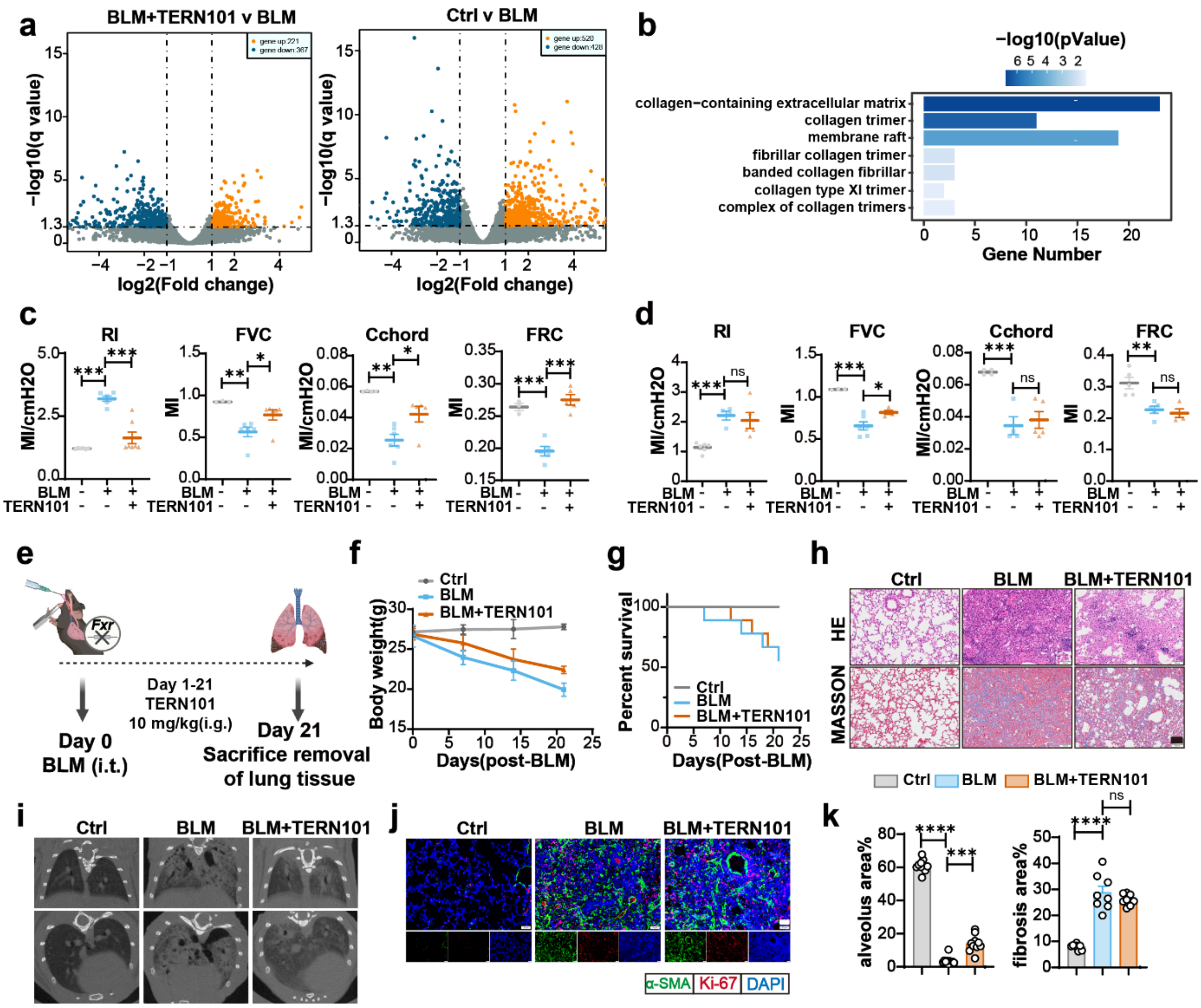

**Supplementary Fig. 9. Pharmacological activation of FXR attenuates pulmonary fibrosis progression in mice.** (**a**) A volcano plot of RNA-seq. (**b**) Pathway enrichment analysis showing BLM-upregulated pathways inhibited by TERN101. (**c**) pulmonary function in WT mice after BLM challenge with or without TERN101 treatment in a therapeutic model (n = 5 mice per group). (**d**) pulmonary function in *Fxr* $^{-/-}$ mice after BLM challenge with or without TERN101 treatment (n = 5 mice per group). (**e**) Schematic of the BLM-induced pulmonary fibrosis model in *Fxr* $^{-/-}$ mice. (**f-g**) Body weight changes(**f**) and survival curves (**g**) in *Fxr* $^{-/-}$ mice after BLM challenge with or without TERN101 treatment (10 mg/kg). (n = 5 mice per group). (**h, i and k)** Histopathological assessment of pulmonary fibrosis in BLM-challenged mice by H&E staining, Masson's trichrome (**h, k**), and micro-CT images(**i**). Scale bars: 100 µm (H&E/Masson), 1 mm (micro-CT). (**j**) Representative immunofluorescence images of lung tissue (Red: Ki67; Green: α-SMA; Blue: DAPI) (Scar bar: 50 µm). The data were assessed by two-tailed Student's t test. *$p < 0.05$; **$p < 0.01$; ***$p < 0.001$; ****$p < 0.0001$; ns, not significant. Data were presented as mean ± SEM.

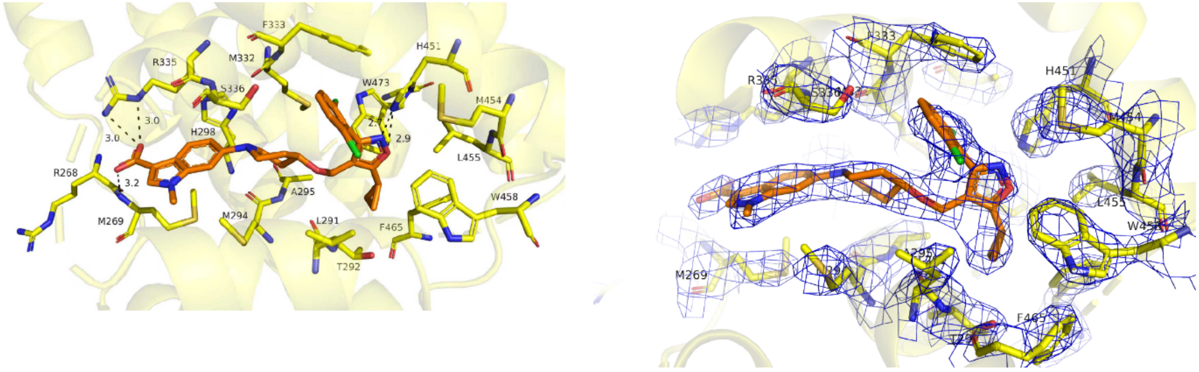

**Supplementary Fig. 10.** Crystal structure of FXR LBD bound with the TERN101 and SRC1-3 peptide (PDB ID: 9WF5), FXR was shown as yellow cartoon, TERN101 and the residues from the binding pocket were shown as orange and yellow sticks, respectively. The 2Fo-Fc electron density map (contour level 1.0 σ) of TERN101 and residues from the binding pocket were shown as blue mesh.

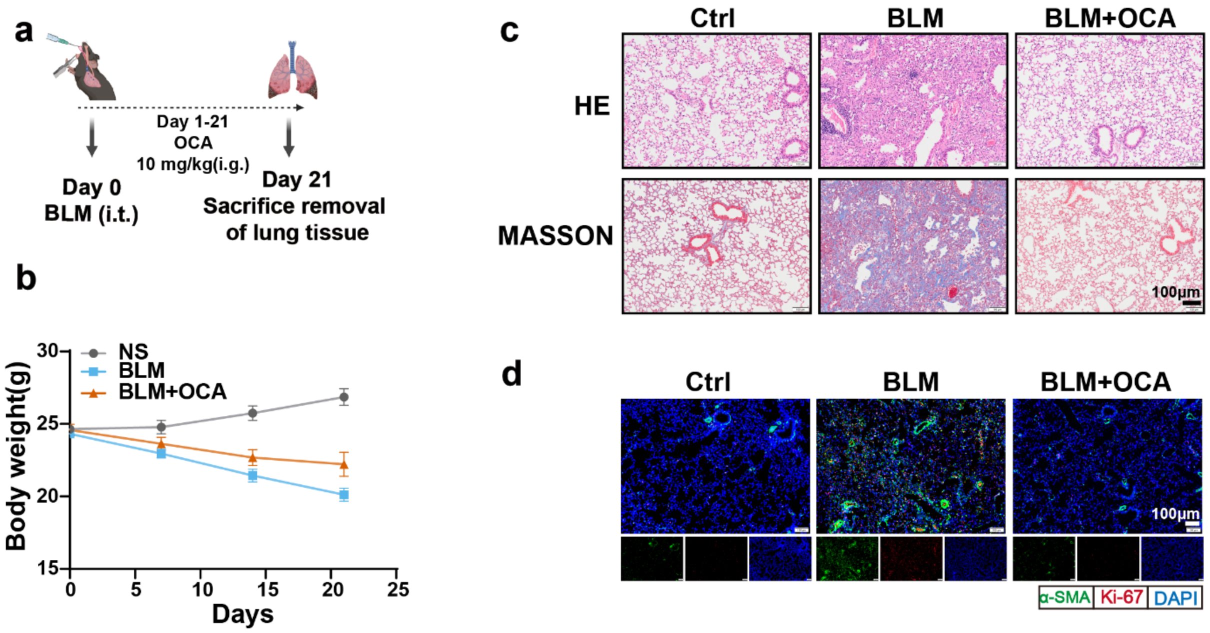

**Supplementary Fig. 11. FXR agonist OCA attenuates pulmonary fibrosis progression in mice.** (**a**) A Schematic figure of the preventive study. (**b**) Body weight changes in WT mice after BLM challenge with or without OCA treatment (n = 6 mice per group). (**c**) Histopathological assessment of pulmonary fibrosis in BLM-challenged mice by H&E staining, Masson's trichrome(Scar bar: 100 μm). (**d**) Representative immunofluorescence images of lung tissue (Red: ki67; Green: α-SMA; Blue: DAPI) (Scar bar: 100 μm). Data were presented as mean ± SEM.

#### S2: Supplementary Tables

#### Supplementary Table 1 | X-ray data collection and refinement statistics

| | FXR in complex with TERN-101 |
|---|---|
| **Data collection** | SSRF-BL19U1 |
| Space group | $P\ 2\ 2_1\ 2_1$ |
| Cell dimensions | |
| *a*, *b*, *c* (Å) | 37.35, 98.42, 129.66 |
| *α*, *β*, *γ* (°) | 90, 90, 90 |
| Resolution | 78.39 – 2.95<br>(2.95 – 3.13) |
| $R_{pim}$ | 0.030 (0.129) |
| I / σ(I) | 17.8 (6.0) |
| *CC*1/2 in highest shell | 0.957 |
| Completeness (%) | 100 (100) |
| Redundancy | 11.7 (12.0) |
| **Refinement** | |
| Resolution (Å) | 2.95 |
| No. of unique reflections | 10694 |
| $R_{work}/R_{free}$ (%) | 25.72/27.47 |
| No. of atoms | |
| Protein | 3689 |
| *B*-factors | |
| Protein | 57.35 |
| Ligand | 62.85 |
| RMSD | |
| Bond lengths (Å) | 0.008 |
| Bond angles (°) | 1.252 |
| Ramachandran favored (%) | 98.4 |
| Ramachandran outliers (%) | 0 |
| PDB accession code | 9WF5 |

**Values in parentheses indicate highest resolution shell.*

**Supplementary Table 2 | Primer sequences for RT-qPCR**

| Gene | Primer pairs（5′-3′） |
|---|---|
| Hum-GAPDH(F) | CAGGAGGCATTGCTGATGAT |
| Hum-GAPDH(R) | GAAGGCTGGGGCTCATTT |
| Hum-COL1A1 (F) | GAGGGCCAAGACGAAGACATC |
| Hum-COL1A1 (R) | CAGATCACGTCATCGCACAAC |
| Hum-FN1 (F) | GAGAATAAGCTGTACCATCGCAA |
| Hum- FN1 (R) | CGACCACATAGGAAGTCCCAG |
| Hum-α-SMA (F) | GACGCTGAAGTATCCGATAGAACACG |
| Hum-α-SMA (R) | CACCATCTCCAGAGTCCAGCACAAT |
| M-GAPDH-(F) | GCACAGTCAAGGCCGAGAAT |
| M-GAPDH-(R) | GCCTTCTCCATGGTGGTGAA |
| M-Col1a1-(F) | TAAGGGTCCCCAATGGTGAGA |
| M-Col1a1-(R) | GGGTCCCTCGACTCCTACAT |
| M-α-SMA-(F) | GGACGTACAACTGGTATTGTGC |
| M-α-SMA-(R) | TCGGCAGTAGTCACGAAGGA |
| M-Fn1 (F) | GATGTCCGAACAGCTATTTACCA |
| M-Fn1 (R) | CCTTGCGACTTCAGCCACT |

**Supplementary Table 3 | shRNA sequences**

| Gene | Primer（5´-3´） |
|---|---|
| ShPPP1CB-1# | GAGGAAACCATGAGTGTGCTA |
| ShPPP1CB-2# | TTTCTGAATCGTCATGATTTA |
| ShFXR-1# | GCCTCTGGATACCACTATAAT |
| ShFXR-2# | GCCTGACTGAATTACGGACAT |
| ShSMAD3 | GCCTCAGTGACAGCGCGCTATTT |
| ShSMAD2 | CAAGTACTCCTTGCTGGATTG |

**Supplementary Table 4 | Clinical features of patients with IPF**

| ID | Age | Sex |
|---|---|---|
| 1 | 42 | Male |
| 2 | 63 | Male |
| 3 | 63 | Male |
| 4 | 54 | Female |
| 5 | 63 | Male |
| 6 | 62 | Male |
| 7 | 71 | Male |
| 8 | 65 | Male |
| 9 | 54 | Male |
| 10 | 57 | Male |
